# Novel biomarkers in mutation-specific gene profiles highlight the involvement of COL9A3 in cancer cell plasticity in uveal melanoma

**DOI:** 10.1101/2025.04.24.650445

**Authors:** Q.C.C. van den Bosch, R. M. Verdijk, H. Fadlelseed, T.P.P. van den Bosch, S. Owens, S. Kennedy, E. Kilic, E. Brosens

**Affiliations:** Department of Ophthalmology, Erasmus MC, Rotterdam, The Netherlands; Department of Clinical Genetics, Erasmus MC, Rotterdam, The Netherlands; Department of Pathology, Erasmus MC, Rotterdam, The Netherlands; National Ophthalmic Pathology Laboratory, Royal Victoria Eye and Ear Hospital and Research Foundation, Dublin, Ireland

## Abstract

UM is a deadly ocular malignancy with well-described genetic alterations that predict disease outcome. However, our current understanding of the biological underpinnings of high-risk uveal melanoma progression remains relatively limited. On the basis of RNA expression profiles, we identified 12 novel biomarkers associated with high-risk UM, with protein expression validation of the 2 top markers RBFOX2 and COL9A3. Moreover, we investigated the functional contribution of COL9A3 via its overexpression in low-risk and high-risk UM cell lines. Our data suggest that COL9A3 enhances cell motility and cell proliferation and alters morphology specifically in high-risk UM cells. Furthermore, zebrafish xenograft studies revealed improved cell dissemination in a low-risk UM cell line upon overexpression of COL9A3, whereas a high-risk UM cell line remained under wild-type conditions. Interestingly, RNA sequencing revealed a high-stress profile in high-risk UM cell lines overexpressing COL9A3 and suggested the upregulation of plasticity markers, such as CD44, Nestin, EZH2, ABCB5, PAX3 and CD166, as a coping mechanism. These markers are known as differentiation markers during the development of melanocyte biogenesis and are implicated in plasticity in other high-risk cancers. The relationship between COL9A3 and plasticity was validated by multiplex immunohistochemistry of FFPE tissue, which demonstrated colocalization of COL9A3, CD44 and Nestin. *BAP1*^mut^-UM samples presented higher levels of COL9A3, CD44 and Nestin expression than *EIF1AX*^mut^- or *SF3B1*^mut^ UM samples did. A positive correlation between COL9A3 and Nestin expression in triple-positive cells, regardless of the UM subtype, was observed. Together, our data suggest that COL9A3 is a novel high-risk biomarker that may be involved in UM cell plasticity and is most prevalent in *BAP1*^*mut*^ UM.

## 1. Introduction

Uveal melanoma (UM) is the most common primary intraocular malignancy, with established driver and secondary mutations that predict disease progression[1]. UM is a deadly malignancy with a high probability of metastatic disease[2], and clinical care is currently lacking treatment options for this disease. Despite excellent control of primary tumors, once metastatic disease occurs, only a subset of patients are eligible for immunotherapy. The main reason for eligibility is HLA compatibility with respect to the immunotherapeutic tebentafusp[3]. For over 40 years, in vivo models have failed to improve therapeutic options for metastatic disease, highlighting the need for a deeper understanding of UM biology[4].

The key elements in tumor progression are proliferation at the primary site followed by dedifferentiation toward a more stem-like cell state to allow for self-renewal and migratory and invasive behavior [5]. This is a crucial process necessary for metastatic disease and is generally understudied in UM. UM is a well-defined genetically characterized tumor, with driver mutations typically arising in *GNAQ or GNA11* or at a lower frequency in *PLBC4 or CYSLTR2*. Driver mutations are found in nevi[6, 7], suggesting that for UM to flourish, secondary mutations are key to unlocking phenotypic plasticity. The core secondary mutations in UM are generally point mutations in *EIF1AX*, hotspot mutations in R625 in *SF3B1* or loss-of-function mutations in *BAP1*[8, 9]. Secondary mutations are strongly associated with disease progression, and UM patients are subdivided into low-risk (*EIF1AX*^*mut*^), intermediate-risk (*SF3B1*^*mut*^), and high-risk (*BAP1*^*mut*^) patients[10]. As secondary mutations are generally mutually exclusive and result in distinct disease progression, we hypothesize that key elements involved in early tumor progression and local invasion are dependent on mutational subtype. This hypothesis is supported by our previous work illustrating methylation-specific profiles that cluster to their representative secondary mutations. Additionally, metastatic primary UM and matched UM metastases present similar epigenetic profiles, suggesting that metastatic capacity develops early in primary UM. However, epigenetic alterations unique to metastasis are patient-specific[11].

Cancer is a heterogeneous disease in which cells are transcriptionally in different cell states[12]. We hypothesize that the dedifferentiation of cancer cells toward a more stem-cell-like state is dependent on the cell of origin, where intermediate states toward embryonic-like pluripotency might mimic early differentiation subtypes of the cell of origin. To identify regulatory genes involved in the early development of uveal melanocytes, we previously identified a list of genes involved in the early development of ocular pigmentation using zebrafish larvae as a model[13]. Recently, comparisons between normal uveal melanocytes and 2 aggressive PDX models revealed the upregulation of PRAME, EPHA4 and EZH2[14]. Particularly interesting is the upregulation of EZH2, as it has been linked to the development of cancer stem cells[15]. Owing to mutation-specific disease progression, we hypothesize that early uveal melanocytic developmental genes upregulated in high-risk UM could unravel mutation-specific dedifferentiation via functional analysis of novel biomarkers and could subsequently serve as potential therapeutic targets. Here, we evaluated 480 genes implicated in early ocular pigmentation in a mutation-specific fashion in 3 distinct RNA-seq cohorts and validated the protein expression of the top markers in 2 patient cohorts. Ultimately, we carried out functional analysis of the validated biomarker COL9A3 and validated the experimental findings in patient tissue.

## 2. Methods

### 2.1 Discovery of novel biomarkers and identification of altered pathways via RNA-seq

Novel biomarkers were identified by reanalyzing the TCGA uveal melanoma cohort and previously published RNA-seq data from two separate cohorts of the ROMS[16]. The gene expression of previously discovered genes implicated in early vertebrate ocular pigmentation development[13] was studied by comparing the gene expression of 480 genes (Supplementary File S1) between molecularly distinct subgroups.

### 2.2 Total RNA profiling of primary tumors and UM cell lines

Two independent total RNA sequencing experiments were performed. First, in the discovery cohort, we compared BAP1-mutated primary tumors (n=14), SF3B1-mutated primary tumors (n=11) and EIF1AX-mutated primary tumors (n=9) (Supplementary File S2 and S3). Differential expression analysis was performed on primary tumor samples with a known secondary driver mutation or negative BAP1 immunostaining. We replicated the signals from this experiment with an independent second cohort of BAP1-mutated primary tumors (n=7), SF3B1-mutated tumors (n=12) and EIF1AX-mutated primary tumors (n=7)(Supplementary File S4 and S5 [16]). The replication cohort has previously been described, and the data was reanalyzed to match the discovery cohort analysis settings. The replicated differentially expressed molecules in the BAP1, SF3B1 and EIF1AX groups were validated in the corresponding groups of the TCGA cohort (BAP1 (n=23), SF3B1 (n=15) and EIF1AX (n=9) with a known secondary driver mutation(Supplementary File S6 and S7, [17]). These replicated and validated RNA molecules, RNA biomarkers, are considered in downstream analysis. Clinical data and RNA expression values can be found in Supplementary Files. For the UM cell lines 92.1-GFP, 92.1-COL9A3-GFP, MP46-GFP and MP46-COL9A3-GFP, total RNA sequencing experiments were carried out on triplets with variation in their culture passage (at least 1 week of culture between each replicate). Following data normalization and differentially expressed gene expression between the GFP and COL9A3-GFP classes, significantly expressed genes were considered in downstream analysis for pathway analysis. For downstream pathway analysis of UM cell lines, Ingenuity Pathway Analysis (QIAGEN, Aarhus, Denmark) was used. All UM cell lines were analyzed for pathway analysis via the same settings with a minimal fold change of −2 or 2 and a false discovery rate of <p=0.05.

### 2.3 Reanalysis of the mRNA microarray validation cohort

Two datasets were included in this analysis. The first was previously described by Laurent et al. and included 63 uveal melanoma patients[18] (Supplementary Files S8 and S9). Data (GSE22138) were downloaded from the Gene Expression Omnibus (GEO)[19]; the second dataset was previously generated by our group and included 54 patients[20](Supplementary Files S1 and S11). The downloaded raw data files (CELs) were processed via brb-array tools v4 6.2[21] and the Just GCRMA’ (GC content–Robust Multi-Array Average) algorithm to adjust for background intensities. The data were normalized via quantile normalization, variance stabilization and log2 transformation. Genes and markers whose expression differed by at least 1.5-fold from the median in at least 20% of the arrays were retained. In contrast to our in-house dataset, the metadata for GSE22138 did not include BAP1, EIF1AX, or SF3B1 mutation status. To compare datasets, we used the 15-gene expression panel described by Onken et al. [22] to predict the prognostic class. Classification into GEP class 1 or 2 was based on five models: the compound covariate predictor model[23], diagonal linear discriminant analysis[24], nearest neighbor classification[24], support vector machines with a linear kernel[25] and the Bayesian compound covariate predictor. Clinical data and RNA expression values can be found in Supplementary Files.

### 2.4 Patient inclusion criteria for protein analysis

Protein validation of mutation-specific RNA biomarkers was initially tested on a set of patients from the ROMS cohort between 2010 and 2023 (n=52). The criteria for patient inclusion were a known mutation status or chromosomal profile, sufficient FFPE tumor material and an appropriate follow-up time. Following initial immunohistochemical analysis of the 5 top markers, additional samples with similar inclusion criteria from the UM cohort originating from Dublin (n=119) were included. Additionally, all samples were stained for BAP1 and scored by both Rotterdam (QB, RM) and Dublin (HF, SO, SK); only samples with concordant scoring were used in this study.

### 2.5 Immunohistochemistry of PTP4A3, SLC11A2, DHX16, RBFOX2 and COL9A3

Immunohistochemistry was carried out via an automated, validated, and accredited staining system (Ventana Benchmark ULTRA, Ventana Medical Systems, Tucson, AZ, USA) combined with an ultraview Universal Alkaline Phosphatase Red Detection Kit (Ventana Cat. no. 760-501). Following deparaffinization and heat-induced antigen retrieval (CC1 for 32 min), the tissue samples were incubated for 32 min with PTP4A3 (1:3200 Abcam, Cambridge, United Kingdom, Cat.no. ab50276), SLC11A2 (1:1600, Abcam, Cambridge, United Kingdom, Cat.no. ab140977), DHX16 (1:200, ThermoFisher, Bleiswijk, The Netherlands, Cat.no. PA5-62411), RBFOX2 (1:400, ThermoFisher, Bleiswijk, The Netherlands, Cat.no. PA5-52268) or COL9A3 (1:100, Sigma-Aldrich, Amsterdam, The Netherlands Cat. no. HPA040125). Counterstaining was performed with hematoxylin II stain for 12 min and blue coloring reagent for 8 min according to the manufacturer’s instructions (Ventana Benchmark ULTRA, Ventana Medical Systems, Tucson, AZ, USA). For each antibody, an appropriate control tissue sample was used as a positive control.

### 2.6 Chromogenic multiplex immunohistochemistry of COL9A3, SOX10 and PU.1

Multiplex immunohistochemistry was carried out via an automated staining system (Ventana Discovery, Ventana Medical Systems, Tucson, AZ, USA). Following deparaffinization and heat-induced antigen retrieval (CC1, for 32 min), the tissue samples were incubated with COL9A3 (1:100, Sigma-Aldrich, Amsterdam, The Netherlands Cat.no. HPA040125) and detected using a Discovery Purple kit (Ventana Cat.no. 760-229). The COL9A3 antibody was subsequently stripped by incubation with CC2 for 8 min at 100°C. The next antibody used was PU.1 (1:800, Abcam, Cambridge, United Kingdom, Cat.no. ab76543), was incubated for 32 min and detected via a Discovery Yellow kit (Ventana, Cat.no. 760-239). Finally, after an additional antibody stripping step, SOX10 (RTU, Cell Marque, Cat. no. 760--4968) was incubated for 32 min and detected via a Discovery Teal kit (Ventana, Cat. no. 760-247). The samples were not counterstained to prevent interpretation artifacts between blue-colored hematoxylin nuclei and teal-colored SOX10-positive nuclei.

### 2.7 Fluorescent multiplex immunohistochemistry of COL9A3, CD44 and Nestin

Multiplex immunohistochemistry was carried out via an automated staining system (Ventana Discovery, Ventana Medical Systems, Tucson, AZ, USA). Following deparaffinization and heat-induced antigen retrieval (CC1, for 32 min), the tissue samples were incubated with COL9A3 (1:100, Sigma-Aldrich, Amsterdam, The Netherlands Cat.no. HPA040125) and detected using a Discovery Red 610 kit (Ventana Cat.no. 760-245). The COL9A3 antibody was subsequently stripped by incubation with CC2 for 8 min at 100°C. The next antibody, Nestin (1:25600; Novus Biologicals, Cat.no. NB300-265), was incubated for 32 min and detected using a Discovery Cy5 kit (Ventana, Cat.no. 760-238). Finally, after an additional antibody stripping step, CD44 (1:4000, Novus Biologicals, Cat.no. NBP1-47390) was incubated for 32 min and detected via a Discovery FAM kit (Ventana, Cat.no. 760-243). The slides were mounted with mounting medium containing DAPI (Abcam, ab104139) to visualize the nucleus.

### 2.8 Generation of COL9A3-overexpressing UM cell lines

Synthetic COL9A3 cDNA from pGEM-COL9A3 (Sino Biological Inc. Germany, Cat.no. HG13717-G) was amplified and cloned and inserted into pTFORF2004 (Addgene, Watertown MA, USA Cat.no. #143330) by replacing the original cDNA insert via HiFi assembly (New England Biolabs, Ipswich MA, USA, Cat.no. E2621S). Additionally, the pCOL9A3 plasmid was adapted to express eGFP for selection and imaging purposes by removing the puromycin coding region with a CMV-GFP insert from pCMV-GFP (Addgene, Watertown MA, USA, Cat.no. #11153). Lentiviral production was achieved by transfecting HEK293T cells with pMDLg/RRE (Addgene, Watertown MA, USA, Cat.no. #122511), pMD2. G (Addgene, Watertown MA, USA, Cat.no. #12259) and pRSV-Rev (Addgene, Watertown MA, USA, Cat.no. #12253) together with pCOL9A3-eGFP for 48 hours at 37°C in DMEM (Gibco, Cat.no. 41965047) supplemented with 10% fetal calf serum (Gibco, Cat.no. A384001) and 1% nonessential amino acids (Gibco, Cat.no. 11140035). Media containing viral particles were filtered to remove cell debris and sequentially used to transduce 92.1 and MP46 cells. After 24 h, transduction was validated via fluorescence microscopy, and eGFP+ cells were isolated via FACS (Aria) to obtain stable cell lines overexpressing COL9A3 and eGFP.

### 2.9 Cell culture conditions

The melanoma cell lines 92.1 (established at the Leiden University Medical Center, Leiden, The Netherlands[26]), MP38 and MP46 (established at Curie Institute, Paris, France, [27]) were used for this study. The chromosomal aberrations and mutation status and research resource identifiers of all the cell lines can be found in Supplementary Table 1. Melanoma cell lines were authenticated previously via single polymorphism analysis and the AmpFLSTR™ Identifiler™ Plus PCR Amplification Kit (Thermo Fisher, Bleiswijk, The Netherlands), followed by sequencing. All uveal melanocytes were propagated in RPMI supplemented with 20% heat-inactivated fetal calf serum and 1% penicillin-streptomycin; MP38 and MM28 media were supplemented with sodium pyruvate. All the cell lines were incubated at 37°C in a humidified 5% CO2-enriched atmosphere and regularly checked for mycoplasma. For protein analysis of the cell cultures, the cells were centrifuged, fixed in 10% buffered formalin and mixed in 3% low-melting agarose (Thermo Fisher, Bleiswijk, The Netherlands, Cat.no. 16520100). Agar blocks were sequentially processed to obtain FFPE blocks for standardized immunohistochemistry as described above.

### 2.9 Cellular architecture and attachment analysis

A total of 2.5×10^5^ cells were seeded in 48-well plates and imaged on an Olympus IX-70 at the same coordinates at 1, 2, 3, 4, 6, 20 and 48 hours postseeding. A total of 5 wells per cell line were investigated. The obtained images were stacked and analyzed in FIJI to manually count the number of attached and unattached cells. Two additional images per well were subsequently obtained at 48 hours postseeding to analyze the shapes of 40 attached cells per cell line. Cell detection was performed manually in FIJI to measure the cell perimeter, Ferret diameter and circularity. The obtained values were processed in GraphPad V9 (GraphPad Software, Boston, MA, USA) via the Mann-Whitney T test to compare knockout cells with WT cells.

### 2.10 Cell proliferation

Total cell proliferation was quantified via the Opera Phenix high-content screening system (Revvity, Groningen, The Netherlands). A total of 2000 cells per cell line were seeded per well five times in a 96-well plate. Automated detection of the cell surface obtained at 0 hours served as the starting point for quantification. Serial images were obtained every 12 hours for a total of 72 hours. Growth was determined by normalizing the data on the basis of the overall surface coverage at 0 hours and plotting the data in GraphPad V9 to determine the duplication time points of each cell line.

### 2.11 Wound healing assay

Migration capacity was investigated via a 2D invasion model by seeding UM cells and fibroblasts in a culture insert (Ibidi, Gräfelfing, Germany, Cat.no. 80366). After 24 hours, the insert was removed, and the cells were imaged for 24 hours in a temperature-, humidity- and CO_2_-controlled environment via a Nikon WideField fluorescence microscope. Every 10 min, images were captured at multiple locations along the cell-free gap between UM cells and fibroblasts. Time lapse images were analyzed in FIJI[28] to quantify the migratory paths of 16 cells per cell line evaluated. Finally, the acquired data were statistically analyzed in GraphPad V9 via ANOVA with post hoc Dunnett’s test.

### 2.12 Zebrafish xenograft models

Wild-type AB zebrafish were maintained under standard conditions with a 14 hr light and 10 hr dark cycle. In this study, only larval zebrafish (no older than 120 hours post fertilization) were used. The animal experiments were approved by the Animal Experimentation Committee at Erasmus MC, Rotterdam. Zebrafish embryos were raised in E3 medium supplemented with 0.003% phenylthiourea in a Petri dish at 28°C. At 24 hours post fertilization, the medium was changed following dechorionation of the larvae. At 48 hours post fertilization, zebrafish larvae were anesthetized with 0.016% tricane and used for injections. A total of 2.5×10^6^ cells were harvested and stained with 2.5 µM CellTracker CM-Dil dye for 5 minutes at 37°C and then for an additional 15 minutes at 4°C. After staining, the CM-Dil dye was removed via centrifugation. The cells were then washed with PBS and resuspended in 2% PVP-40/PBS. A total of ~200–300 cells were injected into the perivitelline space. Successfully injected larvae were selected 1 hour postinjection under a fluorescence stereomicroscope, placed in E3 medium supplemented with PTU and raised at 34°C. At 3 days postinjection, the xenograft larvae were anesthetized and embedded in 1% low-melting agarose for live-cell imaging.

### 2.13 Confocal microscopy of zebrafish xenografts

Zebrafish xenograft larvae were imaged via a Leica SP5 (Leica Camera, Wetzlar, Germany) under standard conditions (561 nm, 35% laser power with additional bright field images). Tile scans of 3 images were generated to obtain full-body length images for analysis. The number of disseminated cells, the distance of dissemination and the total tumor volume were calculated in FIJI via publicly available scripts described in previous work[29]. The obtained values were then processed in GraphPad Prism V9. Comparisons of the number of detected spots, distance of dissemination and tumor volume between the different cell lines were statistically tested via the Mann-Whitney T test to compare wild-type versus overexpression cells.

## 3. Results

### 3.1 Identification of SF3B1-specific and high-risk-associated RNA biomarkers

RNA-seq data were extracted from the TCGA cohort and reanalyzed together with the RNA-seq datasets ROMS-1 and ROMS-2 from our own cohort via CLC Bio to make the datasets comparable with each other. Initially, the gene expression (TPM values) of 482 genes previously identified [13] were compared between molecular subtypes in the ROMS1 cohort and those validated in the TCGA cohort (Supplementary File S1). Among the 482 genes, we identified 1 upregulated *SF3B1*-specific RNA biomarker, *DHX16* (Figure 1A-D), and multiple high-risk biomarkers (the top 12 genes are plotted in Figure 1E). Most genes are clinically correlated with overall survival on the basis of RNA expression (Supplementary Figure 2). For further investigations, the top-ranking genes were selected on the basis of disease-free survival (DFS) significance and the availability of FFPE validated antibodies for protein validation, resulting in a total of four biomarkers eligible for further investigation (Supplementary Figure 1). First, COL9A3, PTP4A3 and RBFOX2 expression was investigated in the RNA-array dataset GSE22138 and our in-house generated dataset[20]. In general, a similar trend was observed, with the exception of SLC11A2 expression, in the ROMS-array cohort (Supplementary Figure 3). Second, additional validation of the top 4 RNA biomarkers *(COL9A3, PTP4A3, SLC11A2, and RBFOX2*) in the ROMS2 cohort supported previous validation and revealed that they were significantly associated with mutation status or GEP classification (Supplementary Figures 3 and 4). The top 4 RNA biomarkers are correlated with poor prognosis (Figure 1F-I) and were selected for further investigation together with 1 *SF3B1-*specific RNA biomarker (DHX16).

**Figure 1:**
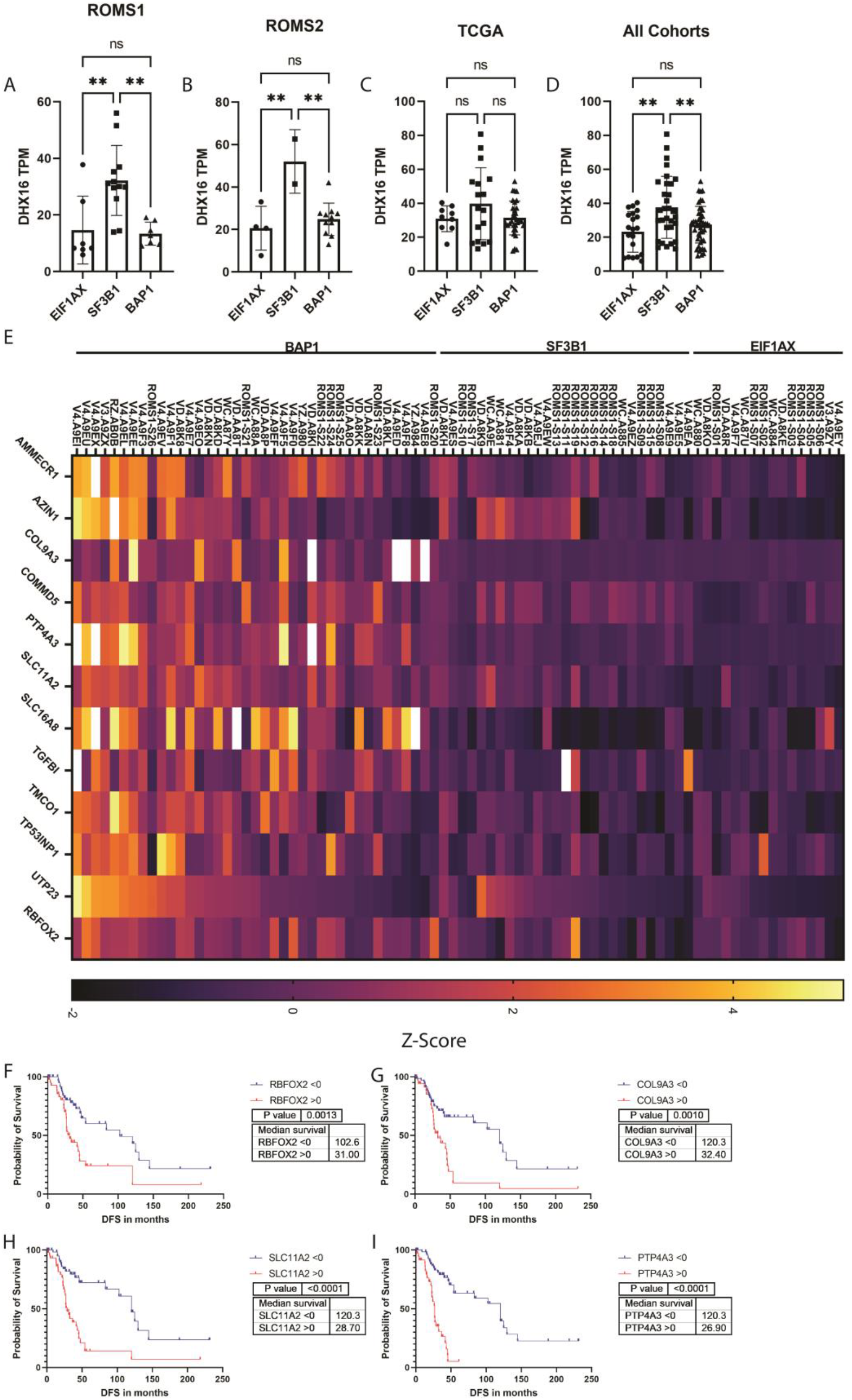
Discovery of novel RNA biomarkers in uveal melanoma. A-D) Expression of the SF3B1-specific marker DHX16 in the ROMS1, ROMS2, TCGA and combined datasets. E) Heatmap of the top markers associated with clinical outcome. F) Kaplan-Meier curve of the combined cohort based on z scores after normalization of the data for RBFOX2. G) Kaplan-Meier curve of the combined cohort based on z scores after normalization of the data for COL9A3. H) Kaplan-Meier curve of the combined cohort based on z scores after normalization of the SLC11A2 data. I) Kaplan-Meier curve of the combined cohort based on z scores after normalization of the data for PTP4A3.

### 3.2 Protein validation of COL9A3, RBFOX2, DHX16, and SLC11A2

Protein expression was determined via immunohistochemistry on FFPE tissue slides via a selection of the ROMS cohort as a discovery set and validated via the Dublin cohort. Initially, protein expression was validated in samples with known RNA data and FFPE available tissue. DHX16, COL9A3, SLC11A2 and RBFOX2 IHC staining confirmed higher protein expression in samples with knownly elevated RNA expression (Supplementary Figure 5), whereas samples stained for PTP4A3 in only 1 out of 3 samples presented elevated protein levels (Supplementary Figure 6). Owing to the RNA-proteinprotein interaction of PTP4A3 and differences between its expression locations (nucleus or cytoplasm), this marker was not evaluated further in the large ROMS and Dublin cohorts. A previous study on PTP4A3 in UM illustrated variation in the staining pattern as well, where strong staining is debatable. Subsequent functional work was performed in OCM cell lines[18], which are derived from cutaneous melanoma, making translatability questionable. The COL9A3 staining pattern suggested the presence of potentially positive macrophages instead of cancer cells themselves because of the presence of foamy-like positive cells (Figure 2A). To validate its expression in either macrophages or cancer cells, we utilized chromogenic multiplex staining with COL9A3 (in red), the macrophage marker PU.1 (in yellow) and the melanocyte marker SOX10 (in blue). Although the COL9A3-positive cells seemed foamy, our multiplex validated expression was only present in cancer cells (Figure 2B). Next, the ROMS cohort was used as a discovery set of 52 FFPE samples to evaluate clinical relevance on the basis of protein expression. In the discovery cohort, RBFOX2 and COL9A3 were the only 2 markers for which elevated protein levels were correlated with poor prognosis (Supplementary Figure 6). Given that COL9A3 is a type of collagen, we wondered if there was any correlation with extracellular matrix patterns such as closed vascular loops. Fisher’s exact test revealed a significant correlation between positive COL9A3 staining and the presence of closed vascular loops in 47 samples (P=0.0031, Supplementary Table 2). DHX16 expression was not correlated with disease-free survival; however, increased expression seems to be elevated in both *BAP1*^*mut*^ and *SF3B1*^*mut*^ UM (Supplementary Figure 7). To validate our initial findings, a larger external cohort from Dublin (n=119) was additionally stained for RBFOX2 and COL9A3. All TMA cores were scored independently by 3 pathologists and 2 researchers with ample experience in ophthalmic histopathology. The obtained scores were compared and discussed to gain concordance between all the samples. The protein level of RBFOX2 was significantly correlated with poor clinical outcome in the ROMS cohort (Figure 2C) and the Dublin cohort (Figure 2D) and was significantly improved after combining both cohorts (Figure 2E). Similarly, COL9A3 protein expression was significantly correlated with clinical outcome in the ROMS cohort (Figure 2F) and the Dublin cohort (Figure 2G) and significantly improved after combining both cohorts (Figure 2H). The samples analyzed additionally presented typical clinical outcomes on the basis of BAP1 staining (Figure 2I).

**Figure 2:**
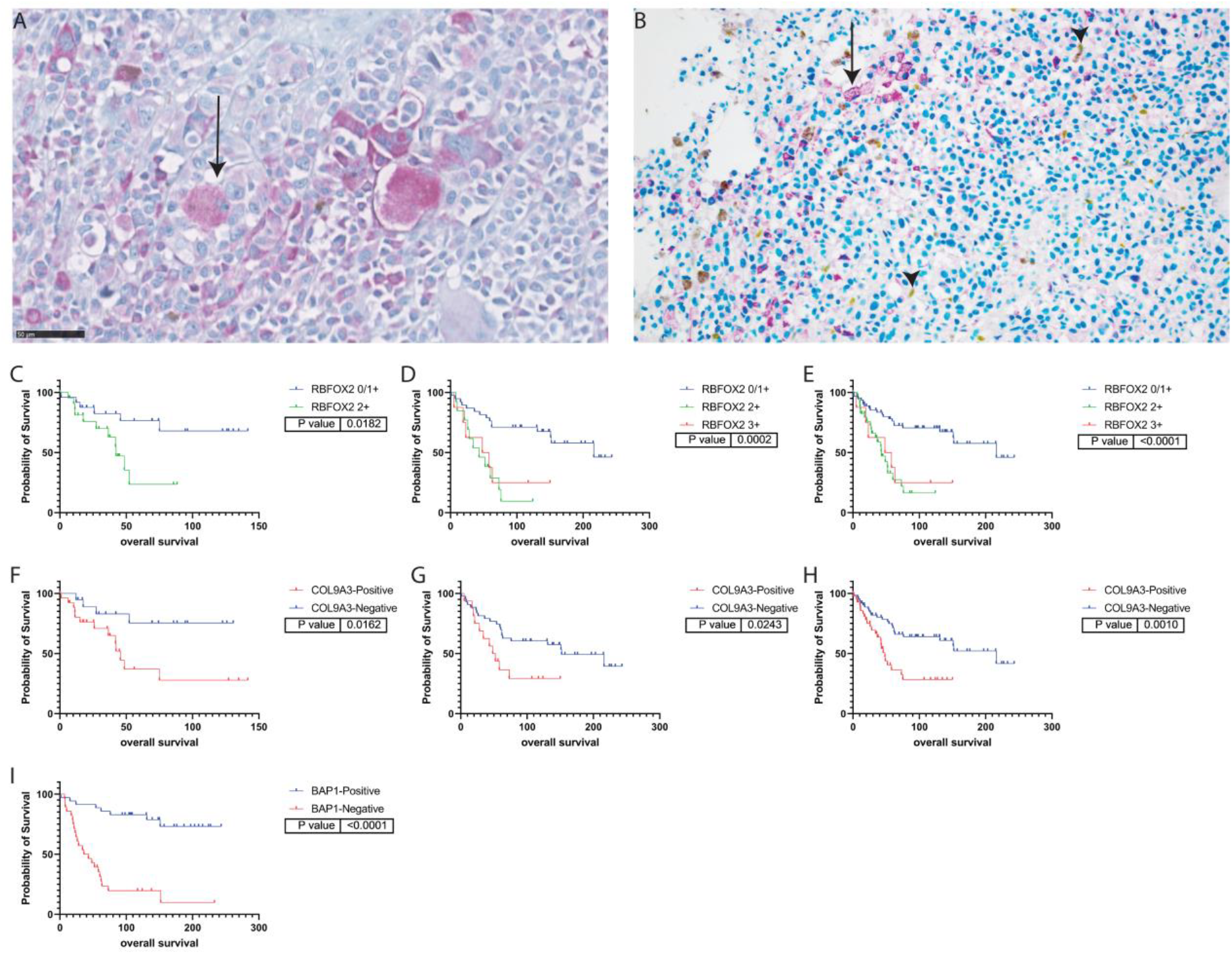
Validation of biomarkers of protein expression. A) Detailed image of COL9A3-positive cells in UM tissue showing a foamy-like cell morphology. B) Chromogenic multiplex of COL9A3 (in red), PU.1 (in yellow) and SOX10 (in blue). The arrows highlight SOX10-positive cells with COL9A3 expression, whereas the arrowheads show PU.1-positive macrophages lacking COL9A3 expression. C-E) Kaplan-Meier curves based on RBFOX2 protein expression in the ROMS, Dublin and combined cohorts. F-H) Kaplan-Meier curves based on COL9A3 protein expression in the ROMS, Dublin and combined cohorts. I) Kaplan-Meier curve based on BAP1 expression.

### 3.4 Validation of COL9A3 overexpression and *in vitro* characterization in UM cell lines

To elucidate the role of COL9A3 in UM progression, stable cell lines overexpressing COL9A3 were established via lentiviral integration. Owing to the strong association between COL9A3 expression and BAP1 status, we investigated its role in both the BAP1 wild-type (92.1 cells) and the BAP1 mutant (MP46) UM cell lines. Following FACS of the GFP-positive cells, immunocytochemical staining was carried out to validate the overexpression of COL9A3 in 92.1 cells (Figure 3A-B) and MP46 cells (Figure 3C-D). The brightfield images were subsequently analyzed for standard morphometrics, which revealed few alterations in 92.1 cells compared with COL9A3-OE cells (Figure 3E-G). Interestingly, overexpression of COL9A3 in 92.1 cells caused significantly slower cell proliferation (best fitted values of a doubling time of 26.07 hours versus 64.70 hours, Figure 3I). On the other hand, overexpression of COL9A3 altered the morphology of MP46 cells, as the OE cells were significantly larger (Figure 3J-L). To evaluate whether COL9A3 influences migration, a wound healing assay revealed a significant improvement in the motility of MP46 cells overexpressing COL9A3 (Figure 3M, Supplementary Figure 8C-D), whereas this improvement was not observed in 92.1 cells (Figure 3L, Supplementary Figure 8A-B). Moreover, OE of COL9A3 significantly improved the doubling time of MP46 cells (best fitted values of a doubling time of 182.8 hours versus 108.2 hours, Figure 3N).

**Figure 3:**
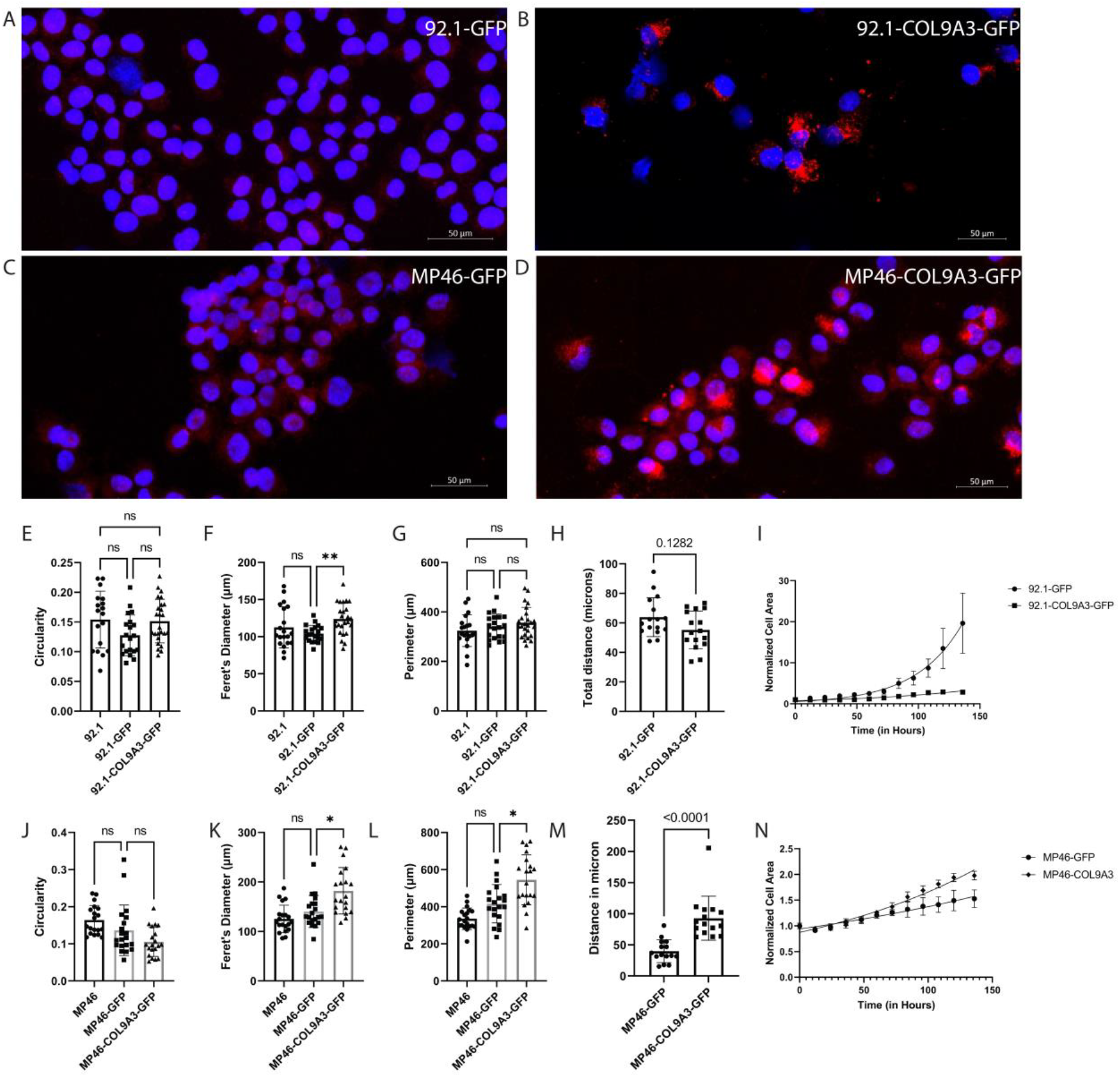
Validation of COL9A3 overexpression and in vitro characterization of low- and high-risk UM cell lines. A) COL9A3 immunocytochemistry of 92.1-GFP cells. B) COL9A3 immunocytochemistry of 92.1-COl9A3-GFP cells. C) COL9A3 immunocytochemistry of MP46-GFP cells. D) COL9A3 immunocytochemistry of MP46-COL9A3-GFP cells. E-G) Morphometric analysis of the circularity, ferret diameter and perimeter of 92.1 cells. H) Quantification of the total distance individual 92.1 cells (n=16 per group) migrated over a time period of 24 hours. I) Cell proliferation assay of 92.1 cells for a total of 144 hours. E-G) Morphometric analysis of the circularity, ferret diameter and perimeter of MP46 cells. H) Quantification of the total distance individual MP46 cells (n=16 per group) migrated over a time period of 24 hours. I) Cell proliferation assay of MP46 cells for a total of 144 hours.

### 3.5 *In vivo* characterization of COL9A3 overexpression in zebrafish xenografts

To further explore the oncogenic ability of COL9A3, we utilized our standardized zebrafish xenograft model[29] to evaluate the tumor burden and cell dissemination capacity in vivo. 92.1-COL9A3-GFP cells did not improve the tumor burden, cell dissemination distance or size of disseminated cell clusters; however, the number of disseminated cells was significantly greater than that of 92.1-GFP cells (Figure 4A-F). Interestingly, MP46-COL9A3-GFP xenografts did not improve the tumor burden, cell dissemination distance, number of disseminated cells or size of disseminated cell clusters (Figure 4G-L). These findings suggest that there could be a threshold to the extent to which COL9A3 influences UM progression, as 92.1-GFP cells naturally have lower COL9A3 expression, whereas MP46-GFP cells have higher expression (Figure 3A, 3C).

**Figure 4:**
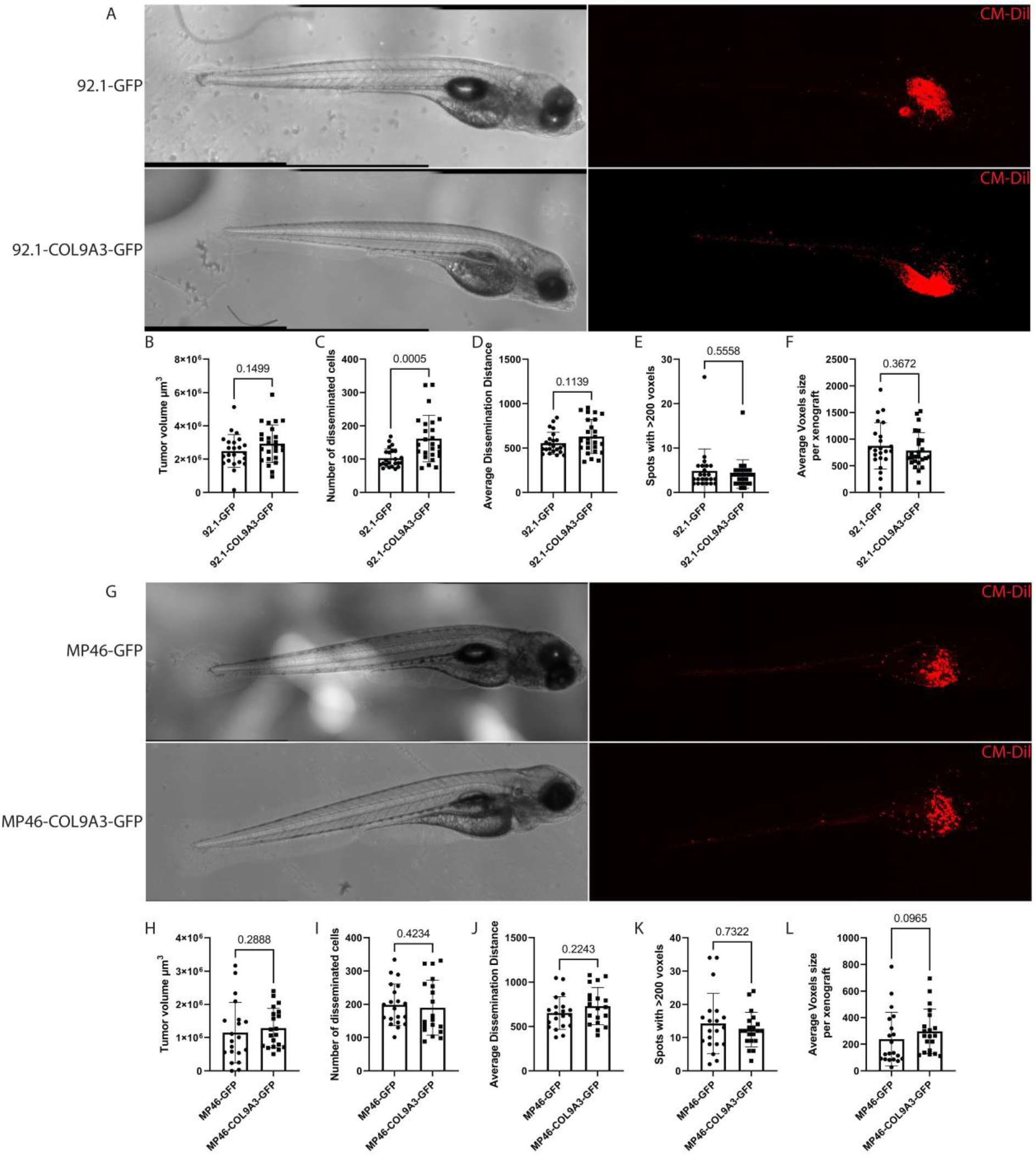
Zebrafish xenograft models of UM cell lines with modified COL9A3 expression. A) Representative image of 92.1-xenografts. B) Total tumor burden in the 92.1-GFP and 92.1-COL9A3-GFP xenograft models. C) Total number of disseminated cells in the 92.1-GFP and 92.1-COL9A3-GFP xenograft models. D) Average distance of disseminated cells from the inoculation site in the 92.1-GFP and 92.1-COL9A3-GFP xenograft models. E) Number of large spots with sizes greater than 200 voxels in the 92.1-GFP and 92.1-COL9A3-GFP xenograft models. F) Overall average voxel size of detected spots per xenograft in 92.1-GFP and 92.1-COL9A3-GFP xenograft models. B) Total tumor burden in the MP46-GFP and MP46-COL9A3-GFP xenograft models. C) Total number of disseminated MP46-GFP and MP46-COL9A3-GFP xenograft model cells. D) Average distance of disseminated cells from the inoculation site in the MP46-GFP and MP46-COL9A3-GFP xenograft models. E) Number of large spots over 200 voxels in the MP46-GFP and MP46-COL9A3-GFP xenograft models. F) Overall average voxel size of detected spots per xenograft in the MP46-GFP and MP46-COL9A3-GFP-xenograft models.

### 3.6 RNA sequencing reveals cellular stress in MP46 cells and suggests that plasticity is a coping mechanism

To understand the influence of COL9A3 in UM, we compared the transcriptomes of stable OE UM cell lines to those of GFP controls. Differential gene expression revealed a strong effect of overexpressing COL9A3 in MP46 cells, with a total of 2895 differentially expressed genes with an FDR p value of >0.05 and a fold change >2 or <-2, whereas the effect on 92.1 cells was minimal, with a total of 91 differentially expressed genes (Figure 5A-B). Pathway analysis via IPA revealed an increase in protein kinase A signaling, the most influential pathway, in 92.1 cells (Figure 5C). However, owing to the large number of differentially expressed genes in MP46-COL9A3-GFP cells, many pathways were altered significantly. In general, MP46-COL9A3-GFP cells exhibit a global loss of RNA translation and processing with additional mitochondrial dysfunction due to the loss of oxidative phosphorylation (Figure 5D). Additionally, significant alterations in cytoskeletal and cytoplasm organization were also observed (Supplementary Figure 9C), which is in line with our morphometric analysis and immunohistochemical staining patterns. This finding suggests high cellular stress in MP46-COL9A3-GFP cells; however, our previous results demonstrated altered morphology and improved in vitro motility and proliferation, whereas comparable in vivo behavior was observed compared with that of MP46-GFP cells. We argued that plasticity markers could provide the mechanism by which MP46-COL9A3-GFP cells overcome cellular stress. An investigation of plasticity markers on the basis of the current literature[30] suggested that MP46-COL9A3-GFP indeed upregulated multiple plasticity markers (Figure 5C-I).

**Figure 5:**
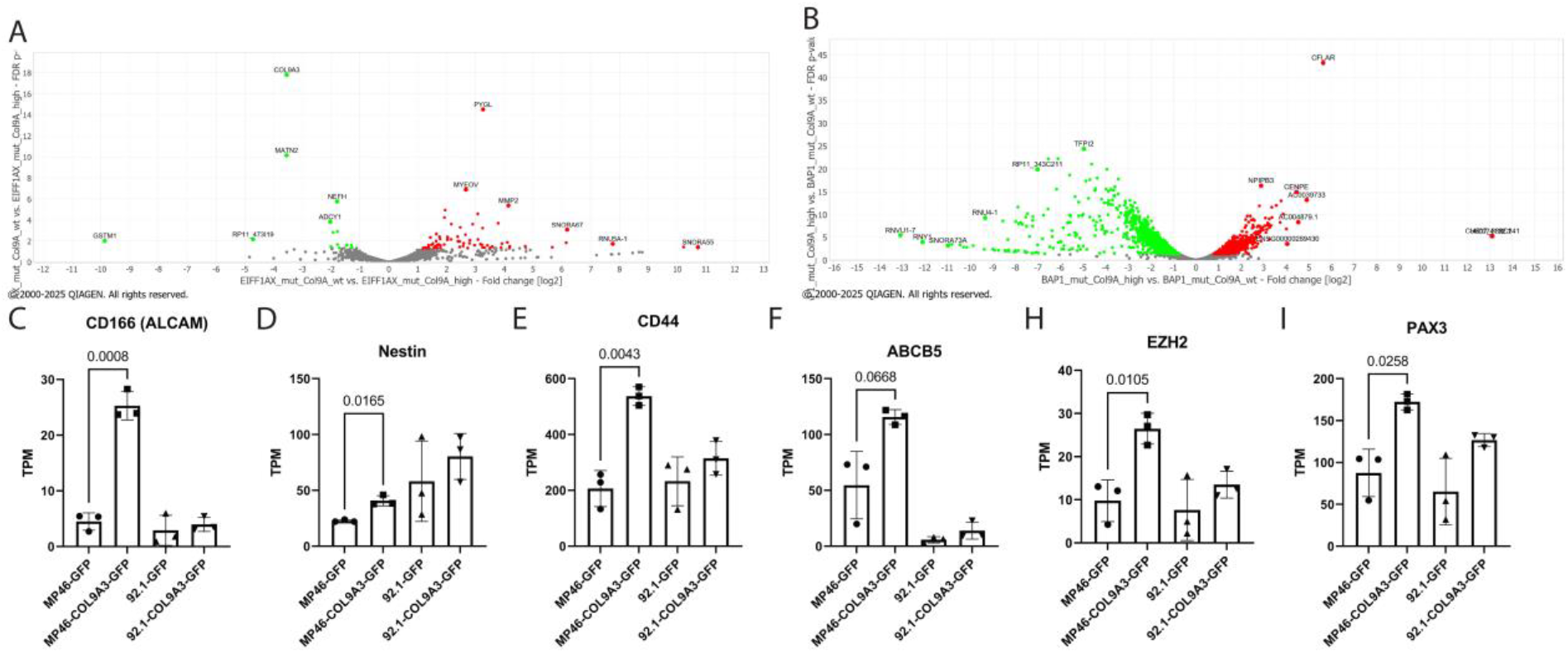
RNA sequencing of COL9A3-modified UM cell lines. A) Volcano plots of differentially expressed genes in 92.1-GFP versus 92.1-COL9A3-GFP cells. B) Volcano plots of differentially expressed genes in MP46-GFP versus MP46-COL9A3-GFP cells. C) Identification of altered pathways via IPA in 92.1-GFP vs 92.1-COL9A3-GFP cells. D) Identification of altered pathways via IPA in MP46-GFP vs MP46-COL9A3-GFP cells. E-J) Expression values of the plasticity markers CD166, Nestin, CD44, ABCB5, EZH2 and PAX3.

### 3.7 Expression of COL9A3 is correlated with plasticity markers in UM patients

Owing to the plasticity association of COL9A3, we wondered whether COL9A3 is involved in dedifferentiation or is a marker for plasticity. Therefore, we investigated the same ROMS-TMA samples used in this study, which allowed the analysis of 36 FFPE samples (N=25 *BAP1*^*mut*^, N=7 *SF3B1*^*mut*^, N=4 *EIF1AX*^*mut*^) via multiplex immunofluorescence staining for COL9A3, CD44 and Nestin (Figure 6A). We developed cell classifiers on the basis of protein expression levels via QuPath to identify single-, double- and triple-positive cells. In general, the BAP1 samples presented the highest percentage of cells positive for COL9A3 alone (Figure 6B) and double positive for COL9A3 with CD44 (Figure 6C) or COL9A3 with Nestin (Figure 6D). Interestingly, the percentage of triple-positive cells did not differ across mutation classes (Figure 6E). Owing to the greater sensitivity of immunofluorescence than chromogenic staining, we investigated protein expression levels at the single-cell level per mutation subtype. The expression of all the markers was highest in the BAP1^mut^ tumor cells (Figure 6F-I). Finally, as our RNA data suggest a relationship between elevated COL9A3 expression and upregulated plasticity markers, we wondered whether triple-positive cells have similar coexpression on the basis of protein quantification. All triple-positive cells were plotted per individual to obtain Spearman’s correlations, which were sequentially plotted together on the basis of their mutational subtype (Figure 6I-J). Interestingly, we observed a generally weak-moderate correlation between COL9A3 and Nestin in all subgroups (Figure 6I), whereas on average, only a weak correlation between COL9A3 and CD44 was observed in SF3B1^mut^ tumor cells (Figure 6J).

**Figure 6:**
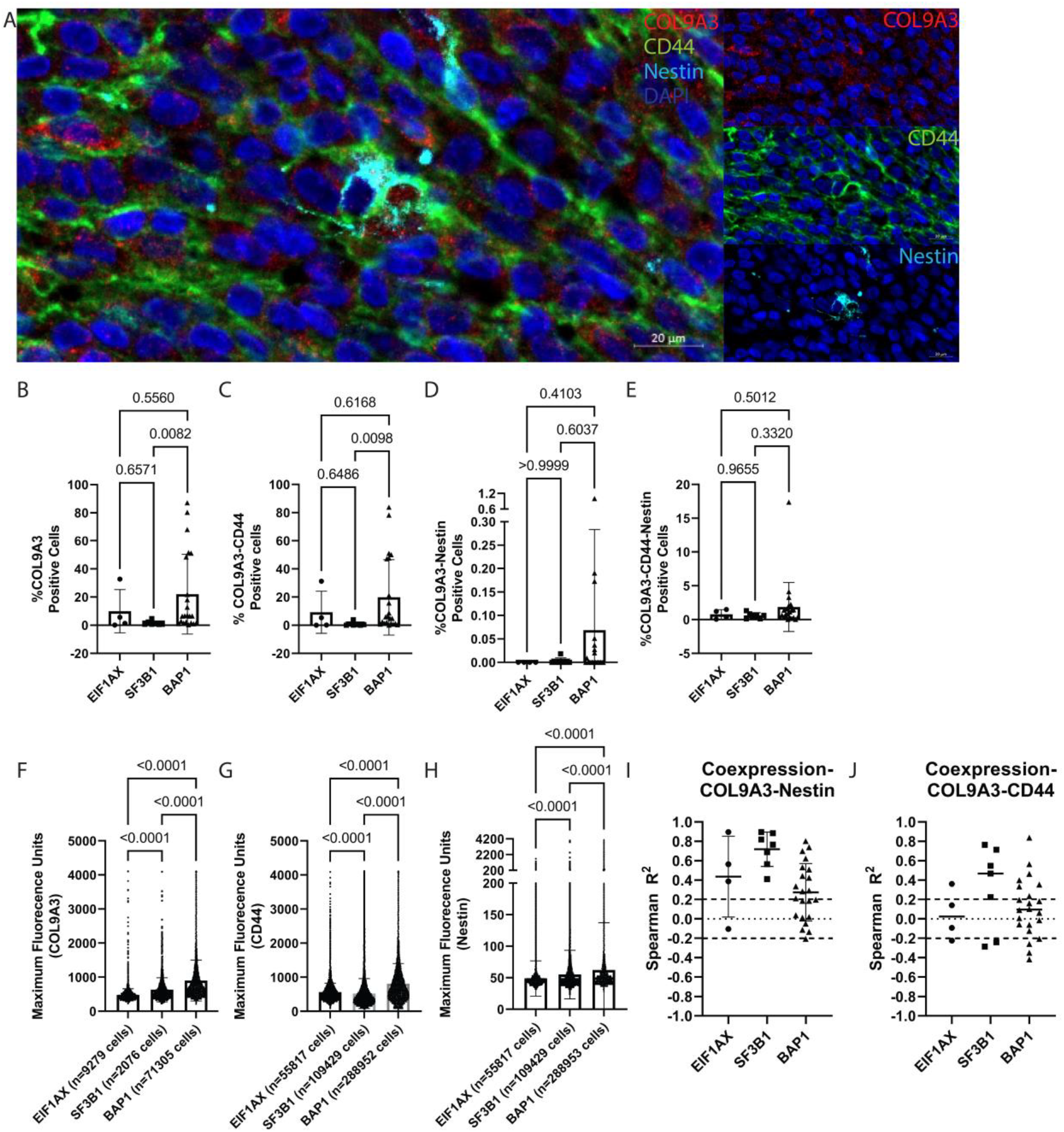
Multiplex immunofluorescence staining of COL9A4, CD44 and Nestin in UM tissue. A) Representative image of COL9A3-, CD44- and Nestin-positive cells; the scale bar represents 20 microns. B) Quantification of positive COL9A3 cells per mutational subgroup on the basis of a percentage of the total number of identified DAPI-positive nuclei. C) Quantification of double-positive COL9A3-CD44 cells per mutational subgroup on the basis of the percentage of the total number of identified DAPI-positive nuclei. D) Quantification of double-positive COL9A3-Nestin-positive cells per mutational subgroup on the basis of the percentage of the total number of identified DAPI-positive nuclei. E) Quantification of triple-positive COL9A3-CD44-Nestin cells per mutational subgroup on the basis of the percentage of the total number of identified DAPI-stained nuclei. F) Maximum fluorescent signal of COL9A3 measured at the individual cell level plotted per mutational subgroup. G) Maximum fluorescent signal of CD44 measured at the individual-cell level plotted per mutational subgroup. H) Maximum fluorescent signal of Nestin measured at the individual cell level plotted per mutational subgroup. I) Spearman correlation of COL9A3 and Nestin expression in triple-positive cells; each value represents an individual tumor. J) Spearman correlation of COL9A3 and CD44 expression in triple-positive cells; each value represents an individual tumor.

## 4. Discussion

In this study, we describe novel biomarkers associated with disease progression in UM at both the RNA and protein levels. From a list of 480 genes, we identified 12 top RNA markers that correlate with poor prognosis. All 12 markers are upregulated in *BAP1*^mut^ UM, suggesting that these could make functional contributions to aggressive behavior, specifically in a BAP1-related context. Furthermore, 2 genes, *RBFOX2* and *COL9A3*, were upregulated at the protein level. To evaluate functional contributions, this study investigated COL9A3 by manipulating UM cell lines and performing in vivo analysis of zebrafish xenografts in a similar fashion as we previously reported that RBFOX2 is involved in metastatic behavior[31]. Our results suggest that COL9A3 is involved in the plasticity of UM cancer cells, which was validated via multiplex immunohistochemistry.

*COL9A3* encodes one of the alpha chains of the type IX collagen heterotrimeric molecule[32], which is part of fibril-associated collagen[33]. Mutations in *COL9A3* have been associated with peripheral vitreoretinal degeneration, retinal detachment[34] and Stickler syndrome[35]. The functional mechanisms of COL9A3 in cancer are currently poorly understood, but its association with high-risk cancer is well established in several cancers. For example, elevated COL9A3 expression is associated with poor prognosis in colorectal cancer[36] [37, 38], gastric cancer[39], osteosarcoma[40], esophageal squamous cell carcinoma[41], salivary adenoid cystic carcinoma[42] and breast cancer[43]. Investigation of the ability of curcumin to suppress LGR5+ colorectal cancer stem cells revealed downregulation of COL9A3. Interestingly, curcumin was able to induce autophagy and inhibit the TFAP2A-mediated ECM pathway, which in turn was shown to be involved in regulating TEAD2 and COL9A3. Moreover, TEAD2 by itself was additionally linked as a regulator of COL9A3[38]. While this study supports our results, where COL9A3 is involved in the plasticity of cancer cells, it also suggests another interesting relationship between COL9A3 and TEAD2. The interplay between COL9A3 and TEAD2 might be interesting to study, as TEAD2 is a known binding partner of YAP[44], which is a crucial gene involved in UM etiology[1, 7]. Additionally, the extracellular matrix is known to be an important factor for stem cells, as it provides structural support and signals to regulate interactions between cells and extracellular matrix components[45]. Although our COL9A3 staining results clearly revealed intracellular expression, there was a correlation between COL9A3 expression and the presence of closed vascular loops, which is a known histological parameter associated with poor prognosis[46]. Therefore, the role of COL9A3 as a structural collagen and interaction protein is of interest because of its role in the plasticity of uveal melanoma cells. In fact, COL9A3 was identified as a downstream extracellular matrix interaction molecule together with COL6A5 in gastric cancer. The role of COL9A3 was identified after studies on USP3 in gastric cancer. USP3 is a deubiquitylase that promotes cell migration and invasion[47], which is mediated by stabilizing COL9A3/COL6A5 in a deubiquitinating-dependent fashion[39]. Furthermore, USP3 is a known chromatin modifier that regulates H2A and H2B[48], which are also targets of BAP1[49]. This link between COL9A3 and a deubiquitylase is particularly interesting, as our study suggests that COL9A3 is specifically relevant for *BAP1*^mut^ UM.

In summary, our results suggest a role for COL9A3 in the plasticity of uveal melanoma cells, which is supported by the current literature with associations with UM-related genes. However, our validation in patient samples remains limited, as only 2 plasticity markers have been evaluated. Further research is needed to establish a direct relationship between plasticity and multiple markers. Additionally, investigations of the interactions of COL9A3 with TEAD2 and deubiquitylation are interesting and could provide novel insights into the plasticity and extracellular matrix interactions of high-risk UM cells. Finally, a deeper understanding of the extracellular matrix and its interactions with UM cancer cells has therapeutic potential for drug development[50].

## 5. Acknowledgments

Some of the data shown here are part of the data generated by the TCGA Research Network: https://www.cancer.gov/tcga.

## Notes

### Competing Interest Statement

The authors have declared no competing interest.

## References

1. Jager, M.J., et al., Uveal melanoma. Nat Rev Dis Primers, 2020. 6(1): p. 24.

2. Kujala, E., T. Mäkitie, and T. Kivelä, Very long-term prognosis of patients with malignant uveal melanoma. Invest Ophthalmol Vis Sci, 2003. 44(11): p. 4651–9.

3. Hassel, J.C., et al., Three-Year Overall Survival with Tebentafusp in Metastatic Uveal Melanoma. N Engl J Med, 2023. 389(24): p. 2256–2266.

4. van den Bosch, Q.C.C., et al., Uveal melanoma modeling in mice and zebrafish. Biochim Biophys Acta Rev Cancer, 2024. 1879(1): p. 189055.

5. Hanahan, D., Hallmarks of Cancer: New Dimensions. Cancer Discov, 2022. 12(1): p. 31–46.

6. Van Raamsdonk, C.D., et al., Frequent somatic mutations of GNAQ in uveal melanoma and blue naevi. Nature, 2009. 457(7229): p. 599–602.

7. Vader, M.J.C., et al., GNAQ and GNA11 mutations and downstream YAP activation in choroidal nevi. Br J Cancer, 2017. 117(6): p. 884–887.

8. Martin, M., et al., Exome sequencing identifies recurrent somatic mutations in EIF1AX and SF3B1 in uveal melanoma with disomy 3. Nat Genet, 2013. 45(8): p. 933–6.

9. Harbour, J.W., et al., Frequent mutation of BAP1 in metastasizing uveal melanomas. Science, 2010. 330(6009): p. 1410–3.

10. Yavuzyigitoglu, S., et al., Uveal Melanomas with SF3B1 Mutations: A Distinct Subclass Associated with Late-Onset Metastases. Ophthalmology, 2016. 123(5): p. 1118–28.

11. Smit, K.N., et al., Genome-wide aberrant methylation in primary metastatic UM and their matched metastases. Sci Rep, 2022. 12(1): p. 42.

12. Barkley, D., et al., Cancer cell states and emergent properties of the dynamic tumor system. Genome Res, 2021. 31(10): p. 1719–1727.

13. van den Bosch, Q.C.C., et al., FOXD1 Is a Transcription Factor Important for Uveal Melanocyte Development and Associated with High-Risk Uveal Melanoma. Cancers (Basel), 2022. 14(15).

14. Gentien, D., et al., Multi-omics comparison of malignant and normal uveal melanocytes reveals molecular features of uveal melanoma. Cell Rep, 2023. 42(9): p. 113132.

15. Chang, C.J. and M.C. Hung, The role of EZH2 in tumour progression. British Journal of Cancer, 2012. 106(2): p. 243–247.

16. Smit, K.N., et al., Aberrant MicroRNA Expression and Its Implications for Uveal Melanoma Metastasis. Cancers (Basel), 2019. 11(6).

17. Robertson, A.G., et al., Comprehensive Molecular Characterization of Muscle-Invasive Bladder Cancer. Cell, 2017. 171(3): p. 540–556 e25.

18. Laurent, C., et al., High PTP4A3 phosphatase expression correlates with metastatic risk in uveal melanoma patients. Cancer Res, 2011. 71(3): p. 666–74.

19. Barrett, T., et al., NCBI GEO: archive for high-throughput functional genomic data. Nucleic Acids Res, 2009. 37(Database issue): p. D885–90.

20. van Gils, W., et al., Gene expression profiling in uveal melanoma: two regions on 3p related to prognosis. Invest Ophthalmol Vis Sci, 2008. 49(10): p. 4254–62.

21. Simon, R., et al., Analysis of gene expression data using BRB-ArrayTools. Cancer Inform, 2007. 3: p. 11-7.

22. Onken, M.D., et al., Collaborative Ocular Oncology Group report number 1: prospective validation of a multi-gene prognostic assay in uveal melanoma. Ophthalmology, 2012. 119(8): p. 1596–603.

23. Radmacher, M.D., L.M. McShane, and R. Simon, A paradigm for class prediction using gene expression profiles. J Comput Biol, 2002. 9(3): p. 505–11.

24. Dudoit, S., F. Jane, and T.P. and Speed, Comparison of Discrimination Methods for the Classification of Tumors Using Gene Expression Data. Journal of the American Statistical Association, 2002. 97(457): p. 77–87.

25. Ramaswamy, S., et al., Multiclass cancer diagnosis using tumor gene expression signatures. Proc Natl Acad Sci U S A, 2001. 98(26): p. 15149–54.

26. De Waard-Siebinga, I., et al., Establishment and characterization of an uveal-melanoma cell line. Int J Cancer, 1995. 62(2): p. 155–61.

27. Amirouchene-Angelozzi, N., et al., Establishment of novel cell lines recapitulating the genetic landscape of uveal melanoma and preclinical validation of mTOR as a therapeutic target. Mol Oncol, 2014. 8(8): p. 1508–20.

28. Schindelin, J., et al., Fiji: an open-source platform for biological-image analysis. Nature Methods, 2012. 9(7): p. 676–682.

29. van den Bosch, Q.C.C., E. Kiliç, and E. Brosens, Uveal Melanoma Zebrafish Xenograft Models Illustrate the Mutation Status-Dependent Effect of Compound Synergism or Antagonism. Investigative Ophthalmology & Visual Science, 2024. 65(10): p. 26–26.

30. Chen, Y.N., Y. Li, and W.B. Wei, Research Progress of Cancer Stem Cells in Uveal Melanoma. Onco Targets Ther, 2020. 13: p. 12243–12252.

31. van den Bosch, Q.C.C., et al., Transcriptional regulators FOXD1 and RBFOX2 contribute to metastatic capacity in <em>BAP1</em><sup><em>mut</em></sup> uveal melanoma. bioRxiv, 2025: p. 2025.04.14.648661.

32. Byron, A., J.D. Humphries, and M.J. Humphries, Defining the extracellular matrix using proteomics. International Journal of Experimental Pathology, 2013. 94(2): p. 75–92.

33. Käpylä, J., et al., The fibril-associated collagen IX provides a novel mechanism for cell adhesion to cartilaginous matrix. J Biol Chem, 2004. 279(49): p. 51677–87.

34. Nash, B.M., et al., Heterozygous COL9A3 variants cause severe peripheral vitreoretinal degeneration and retinal detachment. European Journal of Human Genetics, 2021. 29(5): p. 881–886.

35. Faletra, F., et al., Autosomal recessive Stickler syndrome due to a loss of function mutation in the COL9A3 gene. Am J Med Genet A, 2014. 164A(1): p. 42–7.

36. Yang, X., et al., Potential regulation and prognostic model of colorectal cancer with extracellular matrix genes. Heliyon, 2024. 10(16): p. e36164.

37. Yang, H., et al., An Analysis of the Gene Expression Associated with Lymph Node Metastasis in Colorectal Cancer. Int J Genomics, 2023. 2023: p. 9942663.

38. Mao, X., et al., Curcumin suppresses LGR5(+) colorectal cancer stem cells by inducing autophagy and via repressing TFAP2A-mediated ECM pathway. J Nat Med, 2021. 75(3): p. 590–601.

39. Wu, X., et al., USP3 promotes gastric cancer progression and metastasis by deubiquitination-dependent COL9A3/COL6A5 stabilisation. Cell Death & Disease, 2021. 13(1): p. 10.

40. Wang, Y., et al., Identification of therapeutic targets for osteosarcoma by integrating single-cell RNA sequencing and network pharmacology. Front Pharmacol, 2022. 13: p. 1098800.

41. Liu, M., et al., Profiles of immune cell infiltration and immune-related genes in the tumor microenvironment of esophageal squamous cell carcinoma. BMC Med Genomics, 2021. 14(1): p. 75.

42. Ivanov, S.V., et al., Diagnostic SOX10 gene signatures in salivary adenoid cystic and breast basal-like carcinomas. Br J Cancer, 2013. 109(2): p. 444–51.

43. Lv, X., et al., Identification of potential key genes and pathways predicting pathogenesis and prognosis for triple-negative breast cancer. Cancer Cell International, 2019. 19(1): p. 172.

44. Tian, W., et al., Structural and functional analysis of the YAP-binding domain of human TEAD2. Proc Natl Acad Sci U S A, 2010. 107(16): p. 7293–8.

45. Ragelle, H., et al., Comprehensive proteomic characterization of stem cell-derived extracellular matrices. Biomaterials, 2017. 128: p. 147–159.

46. Kivelä, T., et al., Microvascular loops and networks in uveal melanoma. Can J Ophthalmol, 2004. 39(4): p. 409–21.

47. Wu, X., et al., Ubiquitin-specific protease 3 promotes cell migration and invasion by interacting with and deubiquitinating SUZ12 in gastric cancer. Journal of Experimental & Clinical Cancer Research, 2019. 38(1): p. 277.

48. Nicassio, F., et al., Human USP3 is a chromatin modifier required for S phase progression and genome stability. Curr Biol, 2007. 17(22): p. 1972–7.

49. Carbone, M., et al., BAP1 and cancer. Nat Rev Cancer, 2013. 13(3): p. 153–9.

50. Huang, J., et al., Extracellular matrix and its therapeutic potential for cancer treatment. Signal Transduction and Targeted Therapy, 2021. 6(1): p. 153.

