## Supplementary Data for "Novel biomarkers in mutation-specific gene profiles highlight the involvement of COL9A3 in cancer cell plasticity in uveal melanoma"

**Contents:**

**Supplementary Figure 1: Flow-chart of biomarker discovery used in this study**

**Supplementary Figure 2: Kaplan-Meier survival analysis on 12 Top ranked high-risk associated genes**

**Supplementary Figure 3: Overall-survival and disease-free survival plots of Top 4 high-risk associated genes in re-analyzed RNA-array datasets**

**Supplementary Figure 4: Mutation-specific gene expression of Top 4 selected high-risk associated genes for further experimental investigation**

**Supplementary Figure 5: RNA-Protein validation on tumors with known RNA expression**

**Supplementary Figure 6: RNA-Protein disconcordant of PTP4A3 staining patterns**

**Supplementary Figure 7: Kaplan-Meier survival analysis based on protein expression in the ROMS cohort**

**Supplementary Figure 8: Cell trajectories of individual cells in a wound healing assay for 24 hours.**

**Supplementary Figure 9: Alterations in pathways and cell organization in based on RNA-sequencing data**

**Supplementary Table 1: Genetic information on cell lines used in this study**

**Supplementary Table 2: Fisher’s Exact Test of COL9A3 staining and closed vascular loops**

**Supplementary File: Clinical Data and RNA expression Values**

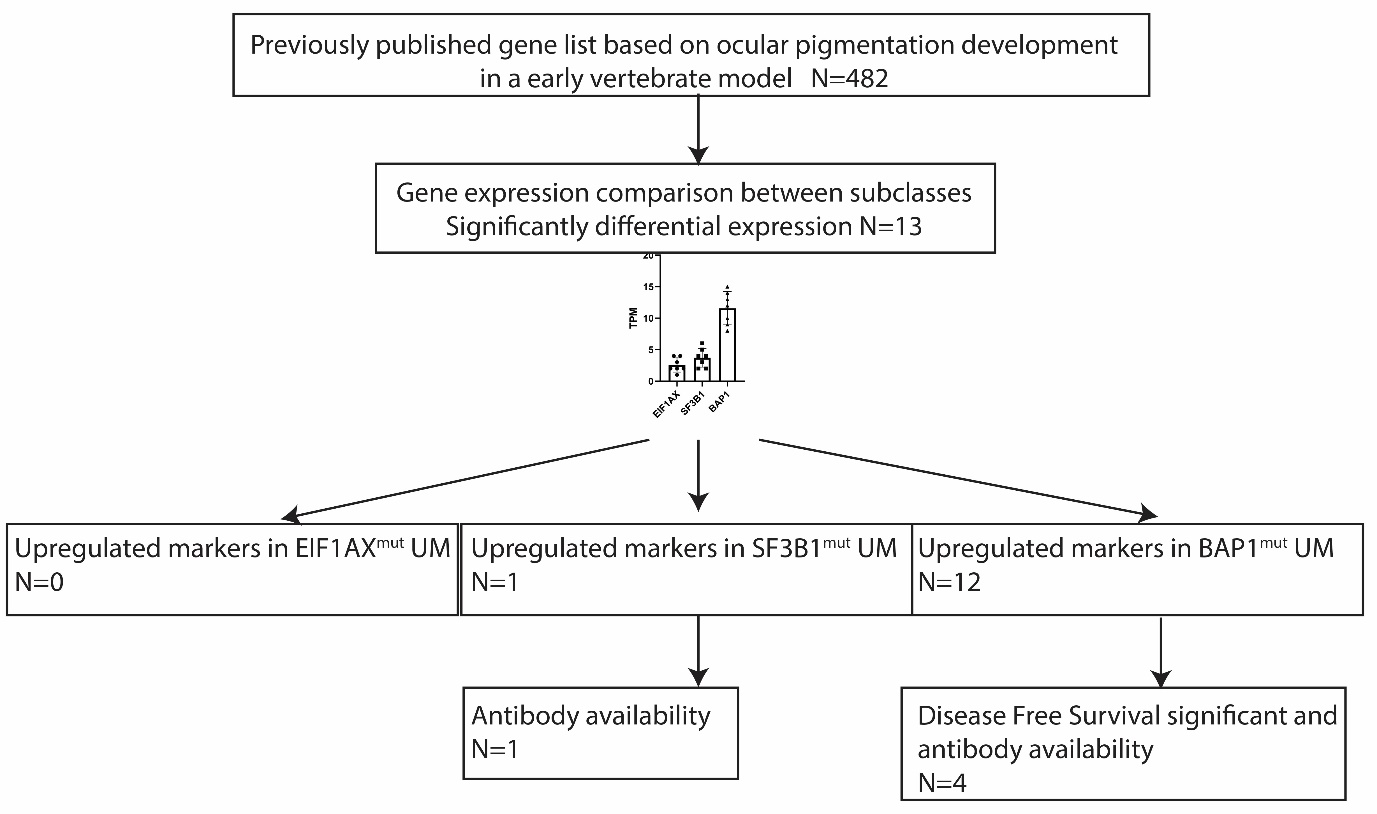

**Supplementary Figure 1: Flow-chart of biomarker discovery used in this study**. From a previously published gene list a total of 482 genes were evaluated for UM subtype expression analysis leading to a total of 14 significant enriched biomarkers. From these 14 markers, 1 marker was specific for SF3B1^mut^ UM whereas 12 were enriched in BAP1^mut^ UM. After availability of FFPE approved antibody selection a total of 5 were subsequently used for immunohistochemistry stainings.

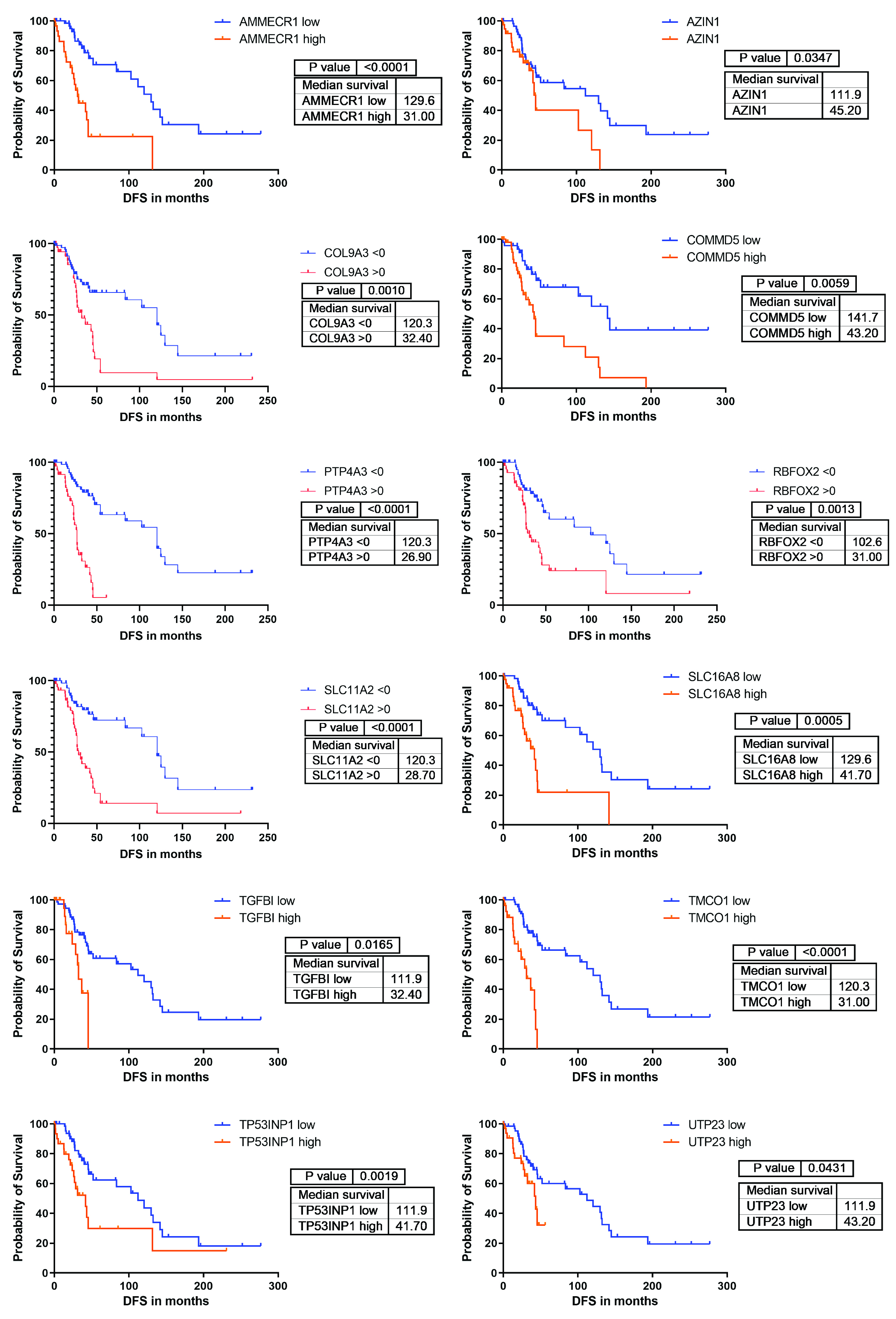

**Supplementary Figure2: Kaplan-Meier survival analysis on 12 Top ranked high-risk associated genes.** Disease free survival based on the RNA expression of AMMERC1, AZIN, COL9A3, COMMD5, PTP4A3, RBFOX2, SLC11A2, SLC16A8, TGFBI, TMCO1, TP53INP and UTP23 of the combined ROMS1, ROMS2 and TCGA cohorts.

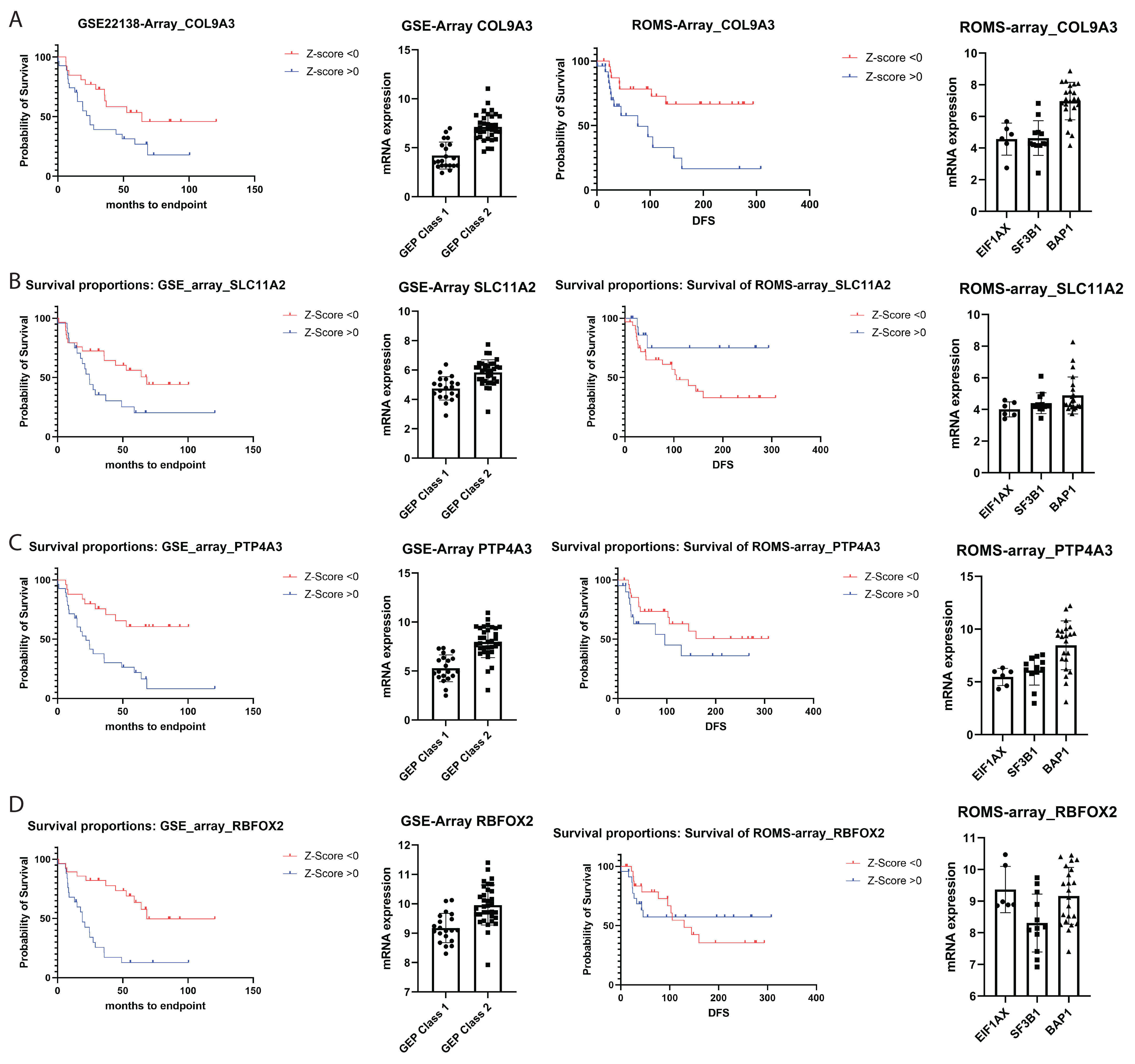

***Supplementary Figure 3: Overall-survival and disease-free survival plots of Top 4 high-risk associated genes in re-analyzed RNA-array datasets.*** *A) Overall survival of GSE22138 and disease free survival of the ROMS-array datasets based on COL9A3 expression normalized to Z-score values. Additionally, expression is linked to either GEP-class or mutational subclass or GSE22138 or ROMS-array samples, respectively. B) Overall survival of GSE22138 and disease free survival of the ROMS-array datasets based on SLC11A2 expression normalized to Z-score values. Additionally, expression is linked to either GEP-class or mutational subclass or GSE22138 or ROMS-array samples, respectively. C) Overall survival of GSE22138 and disease free survival of the ROMS-array datasets based on PTP4A3 expression normalized to Z-score values. Additionally, expression is linked to either GEP-class or mutational subclass or GSE22138 or ROMS-array samples, respectively. D) Overall survival of GSE22138 and disease free survival of the ROMS-array datasets based on RBFOX2 expression normalized to Z-score values. Additionally, expression is linked to either GEP-class or mutational subclass or GSE22138 or ROMS-array samples, respectively.*

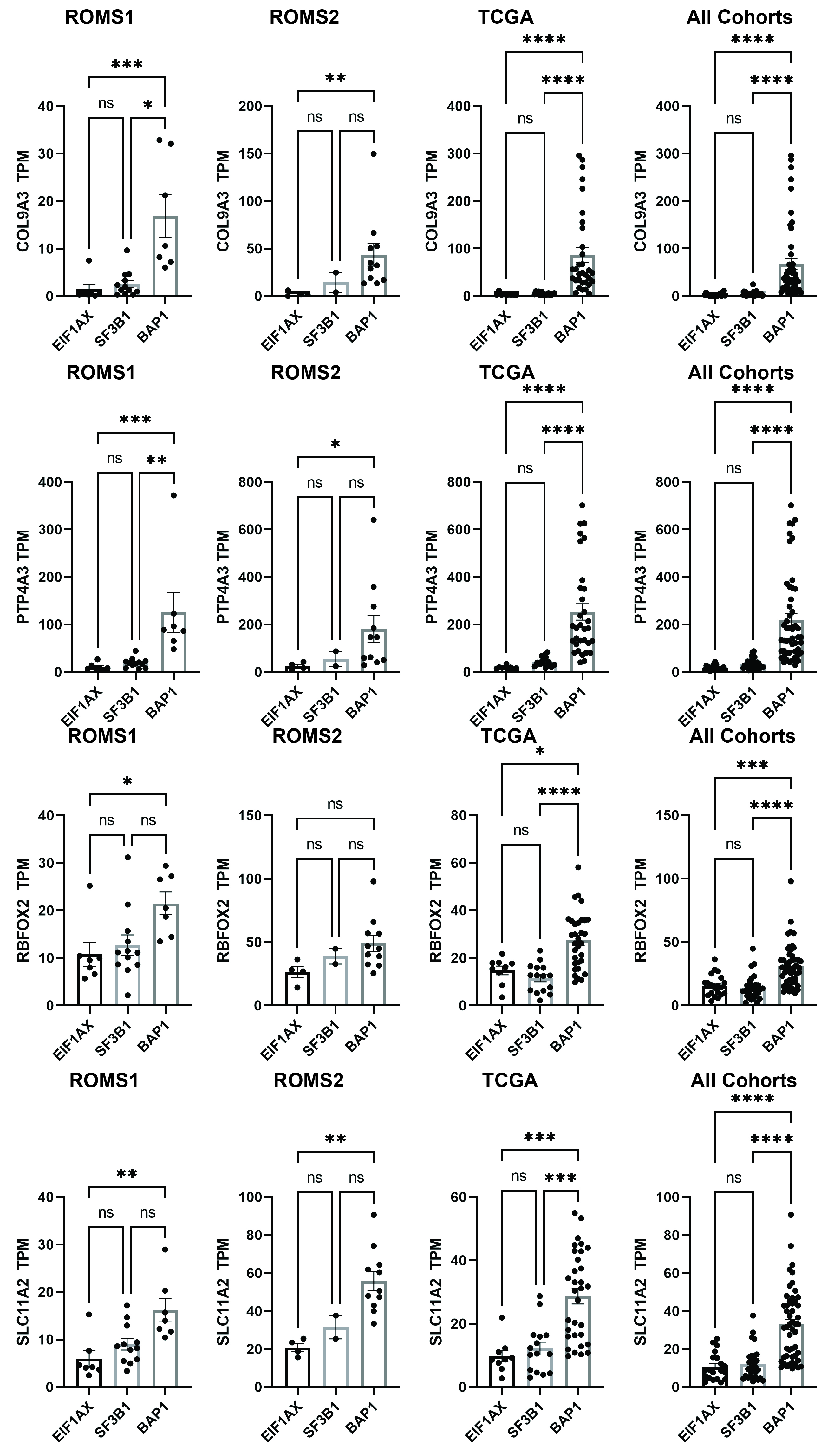

**Supplementary Figure 4: Mutation-specific gene expression of Top 4 selected high-risk associated genes for further experimental investigation.** Biomarkers COL9A3, PTP4A3, RBFOX2 and SLC11A2 that are eligible for antibody staining’s with enriched RNA expression in BAP1^mut^ UM in the ROMS1, ROMS2, TCGA and combined cohorts.

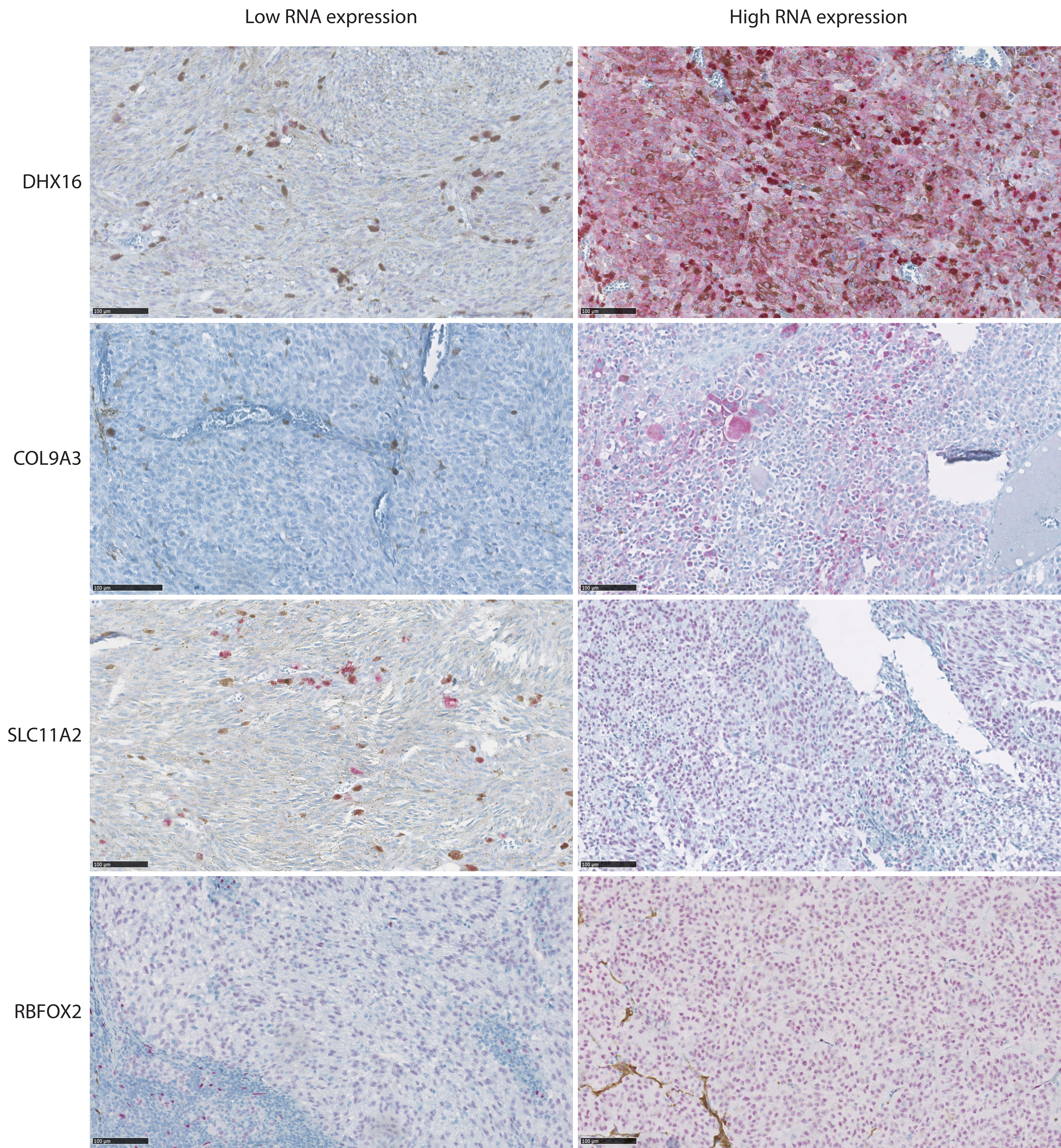

**Supplementary Figure 5: RNA-Protein validation on tumors with known RNA expression.** Immunohistochemistry of DHX16, COL9A3, SLC11A2 and RBFOX2 on UM cases with known RNA expression for RNA-protein validation. Detection of protein expression is shown in red, scale bar represents 100 micron.

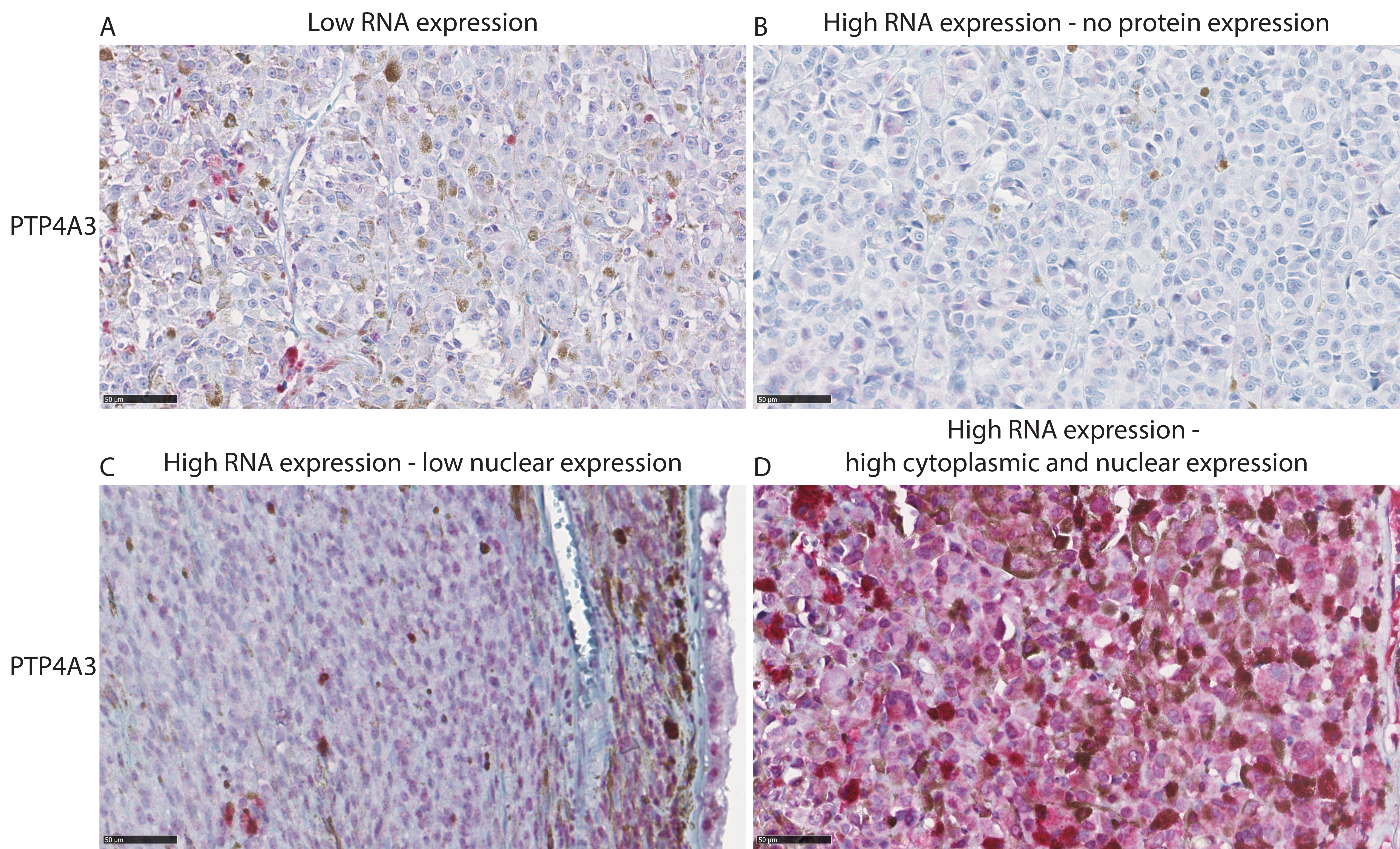

**Supplementary Figure 6: Variation of PTP4A3 staining’s pattern in UM samples with known RNA expression.** RNA-protein validation on UM cases with known PTP4A3 RNA expression illustrate variety in staining patterns. No clear RNA-protein validation could be established due to lack of protein in high-RNA cases, or uncertainty of localization (nuclear, cytoplasmic or both) .Scale bar represents 50 microns.

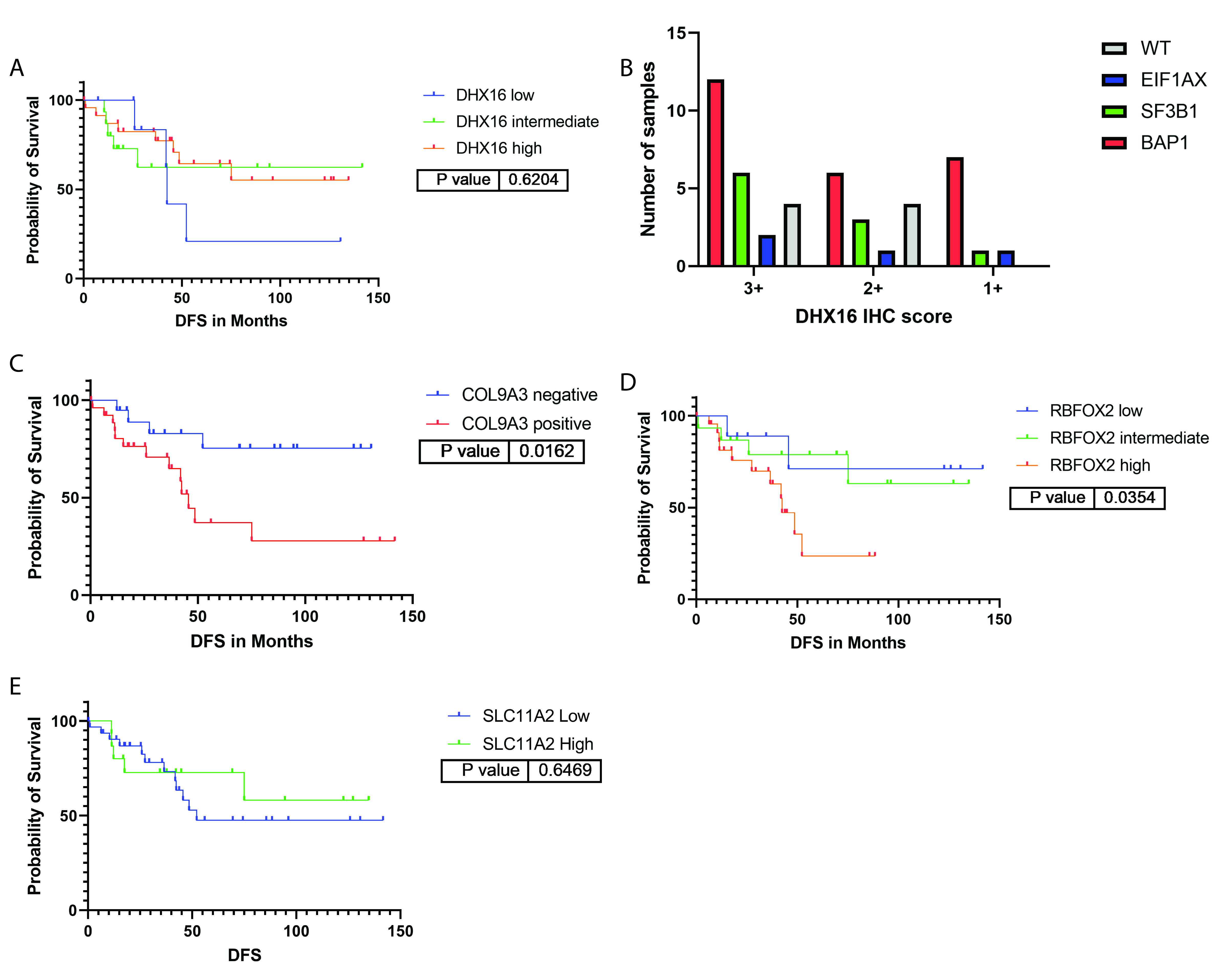

**Supplementary Figure 7: Kaplan-Meier survival analysis based on protein expression in the ROMS cohort.** A) Survival of DHX16 staining on TMA slides lack significant differences in DFS. B) Positive DHX16 cases distributed based on their expression scoring and mutation subtype illustrate increased expression is more common in BAP1 and SF3B1 cases. C) Disease free survival of UM cases based on COL9A3 protein expression. D) Disease free survival of UM cases based on RBFOX2 protein expression. E) Disease free survival of UM cases based on SLC11A2 protein expression.

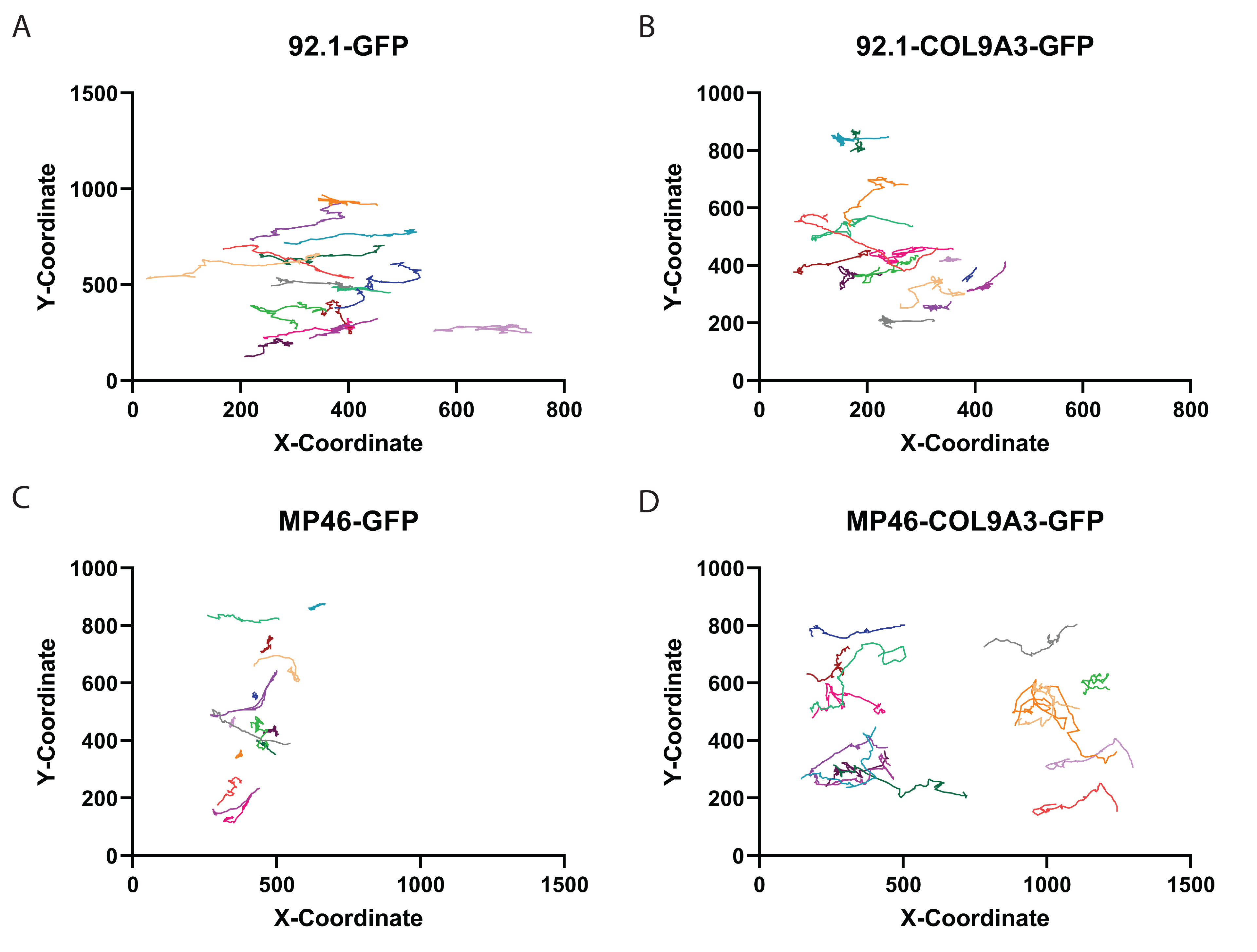

**Supplementary Figure 8: Cell trajectories of individual cells in a wound healing assay for 24 hours**. A) Cell trajectory of 92.1-GFP cells over a time period of 24 hours. B) Cell trajectory of 92.1-COL9A3-GFP cells over a time period of 24 hours A) Cell trajectory of MP46-GFP cells over a time period of 24 hours A) Cell trajectory of MP46-COL9A3-GFP cells over a time period of 24 hours Each color represents a single cell tracked.

**Supplementary Figure 9: Alterations in pathways and cell organization in based on RNA-sequencing data**. A) Pathway enrichment in Ingenuity Pathway Analysis comparing 92.1-COL9A3-GFP cells to 92.1-GFP cells. B) Pathway enrichment in Ingenuity Pathway Analysis comparing MP46-GFP to MP46-COL9A3-GFP cells. C) Alterations of cell organization in MP46-COL9A3-GFP cells compared to MP46-GFP cells.

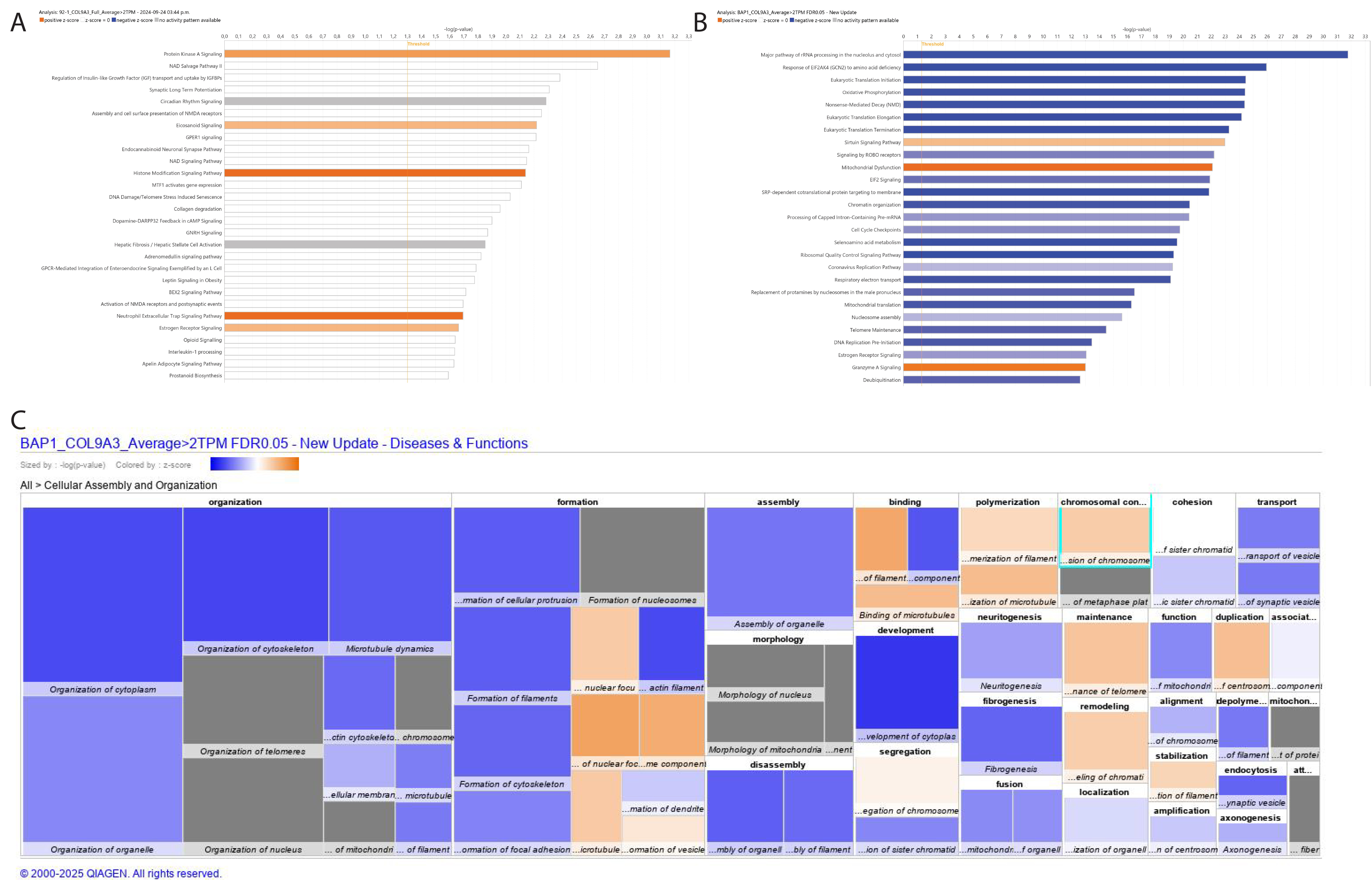

**Supplementary Table 2: Genetic information on cell lines used in this study**

| **Cell line** | **Origin** | **Driver Mutation** | **Secundary Mutation** | **Chr. 3** | **Chr. 6** | **Chr. 8** | **BAP1 protein** | **Ref** | **RRID** |
| --- | --- | --- | --- | --- | --- | --- | --- | --- | --- |
| 92.1 | Primary | GNAQ Q209L c.626A>T | EIF1AX | Disomy 3 | Gain 6p | Gain 8q | Yes | [1] | CVCL_8607 |
|  |  |  | c. 17G/A |  |  |  |  |  |  |
| MP46 | Primary | GNAQ Q209L c.626A>T | WT | Disomy 3, LOH | Gain 6p, Loss 6q | Gain 8q, Loss 8p | No | [2] | CVCL_4D13 |

**Supplementary Table 3: Fisher’s Exact Test of COL9A3 staining and closed vascular loops**

|  | **Cases** | **Presence of closed vascular loops** | **Absence of closed**  **vascular loops** | **Fisher’s Exact Test** |
| --- | --- | --- | --- | --- |
| Positive Staining | 27 | 15 | 12 | 0.0031 |
| Negative staining | 21 | 3 | 19 |  |

**References**

1.	De Waard-Siebinga, I., et al., *Establishment and characterization of an uveal-melanoma cell line.* Int J Cancer, 1995. **62**(2): p. 155-61.

2.	Amirouchene-Angelozzi, N., et al., *Establishment of novel cell lines recapitulating the genetic landscape of uveal melanoma and preclinical validation of mTOR as a therapeutic target.* Mol Oncol, 2014. **8**(8): p. 1508-20.
